# RiboCouple links RNA backbone generation to sequence design through a learned interaction topology

**DOI:** 10.64898/2026.10.15.759394

**Authors:** Zhiqi Ma, Yefeng Zheng

## Abstract

Designing RNAs with prescribed structures is central to RNA engineering. Backbone-first approaches generate geometry before sequence assignment, yet how to transfer relational information to sequence design remains unclear. We introduce RiboCouple, which infers a probabilistic residue-pair state from backbone geometry, recycles pair probabilities through a second pass and transfers them to sequence decoding as soft attention biases. On 550 generated backbones, pair-type decoding achieved joint refolding validity of 45.1% versus 33.5% for a matched topology-free decoder. In another 110-backbone cohort, replacing aligned pair states with statistics-matched donor states reduced validity from 46.4% to 21.4%, demonstrating that decoding depends on backbone-aligned relational information. Before external refolding, pair-confidence ranking enriched validity to 59.6% in the top 248 and transferred to other generators; on a 94-chain temporally held-out native-backbone benchmark, pair-coupled decoding improved secondary-structure agreement. These findings establish backbone-inferred interaction topology as an intermediate representation connecting backbone generation, sequence decoding, and candidate prioritization.

---

Ribonucleic acid (RNA) molecules perform diverse biological functions through their ability to adopt specific and dynamic three-dimensional conformations. Designing RNA molecules with prescribed structures and functions is a central goal of RNA engineering. The dependence of RNA function on structure has motivated computational methods for RNA sequence design. Early approaches used thermodynamic models to optimize sequences for target secondary structures [1, 2], while community-based efforts developed empirical design rules through iterative design and experimental feedback [3]. Subsequent learning-based methods incorporated human design strategies or learned sequence-design policies [4, 5]. Three-dimensional modelling has likewise progressed from physics- and knowledge-based approaches, including fragment assembly, toward learned structure predictors [6–8], with community-wide blind challenges providing benchmarks of progress [9, 10]. Recent generative approaches span sequence-free backbone generation and sequence–structure co-design [11, 12]. In sequence-free backbone-first workflows, the generated geometry does not specify nucleotide identities and therefore requires a downstream sequence-design step.

Despite rapid progress in RNA generation and inverse folding, backbone-first design still lacks a representation-level answer to how generated three-dimensional geometry should be passed to subsequent sequence design. Structure-conditioned inverse-folding models such as gRNAde [13], RDesign [14] and RiboDiffusion [15] design sequences for supplied RNA backbones, whereas RNA-FrameFlow [11] generates sequence-free backbones for downstream sequence design. Joint or conditional frameworks instead couple sequence and geometry within unified generative processes or task-specific conditioning [12, 16–19]. Across these settings, several methods additionally incorporate secondary-structure, base-pair or other pairwise representations into RNA structure or sequence modelling [8, 14, 20–24]. What remains unclear is whether a probabilistic relational state inferred directly from backbone geometry can serve as an explicit interface between backbone generation and sequence decoding—one that can be refined, transferred and examined through controlled perturbations.

RNA provides a useful setting in which to study this problem because its three-dimensional structure is organized by residue-pair interactions. Canonical base pairs form helices, whereas noncanonical interactions, stacking contacts and long-range contacts shape junctions, motifs and higher-order architectures [25, 26]. These interactions constrain the relative geometry of the sugar– phosphate backbone, allowing interaction topology to be inferred probabilistically from residue frames without knowledge of nucleotide sequence. In the sequence-free setting considered here, a generated backbone does not specify the nucleotide at each position or uniquely determine which nucleotide identities are compatible with a given interaction. A probabilistic, typed residue-pair graph therefore provides a natural intermediate: it identifies residues whose nucleotide choices should be coordinated without predetermining either nucleotide. We refer to this backbone-inferred probabilistic residue-pair graph as the interaction topology.

Here we introduce RiboCouple, a backbone-first framework that infers a probabilistic residue-pair topology from generated backbone geometry, recycles binary pair probabilities through a second structural pass and transfers the resulting binary and typed pair signals to nucleotide decoding as soft attention biases (Fig. 1). An SE(3)-equivariant flow-matching trunk generates the RNA backbone while learning residue-level representations. A pair module predicts binary pair probabilities, which are recycled into the pair representation before a second structural pass. The refined representation then produces binary-pair and conditional nucleotide-pair-class logits. Together with the updated residue representations, these logits guide sequence decoding while preserving uncertainty in pair strength and pair class. The inferred interaction topology provides an explicit, backbone-aligned interface for sequence decoding, allowing its contribution to be tested through targeted perturbations at inference.

**Figure 1.**
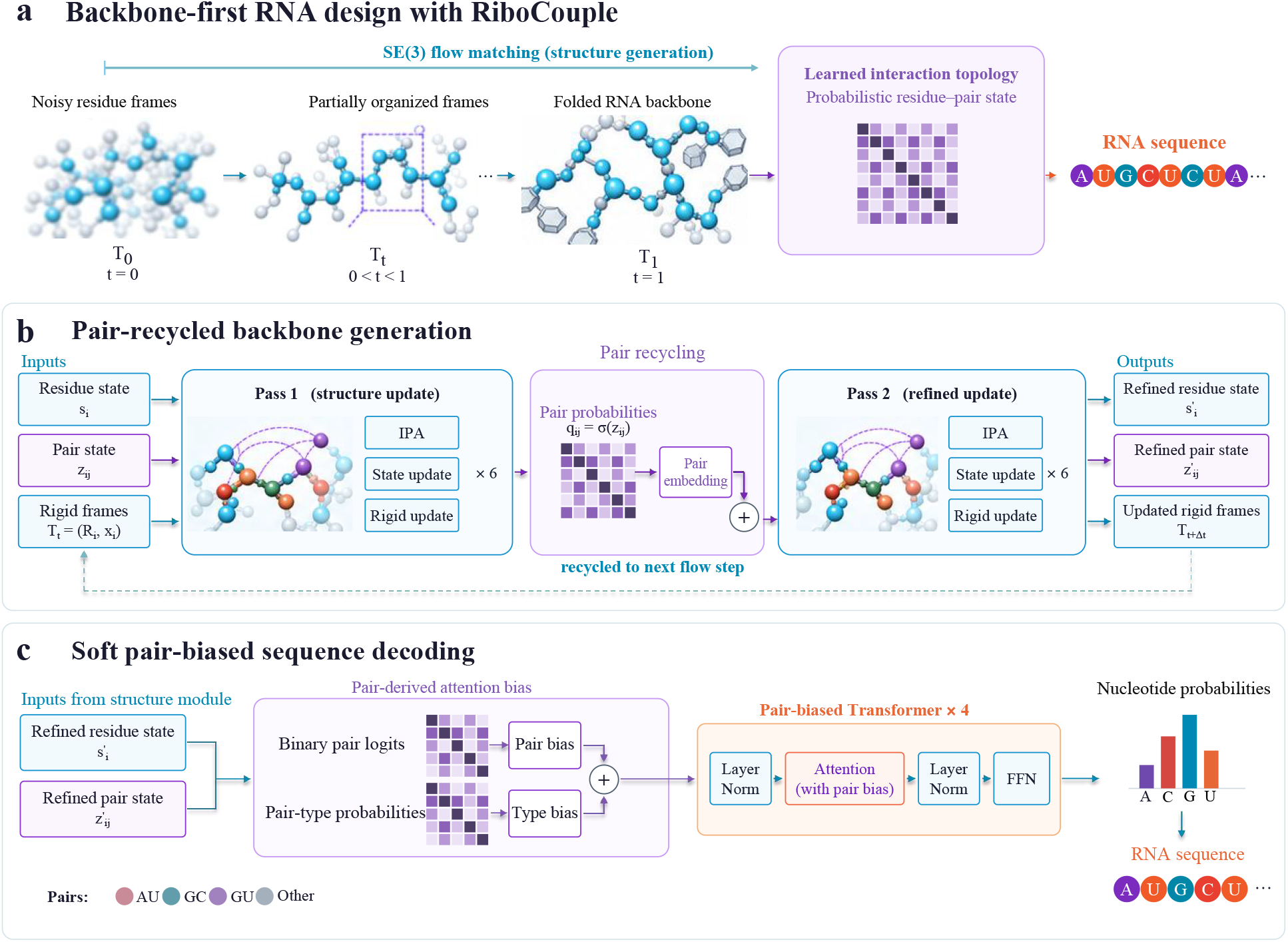
Backbone-first RNA design with learned relational-state transfer. **a**, An SE(3)-equivariant flow-matching model generates RNA backbone geometry while inferring a probabilistic residue-pair state. This learned interaction topology is transferred to sequence decoding. Matrix shading indicates binary pairing probability. **b**, Pair-recycled backbone generation. Residue states (*s*_*i*_), pair states (*z*_*ij*_) and rigid frames (*T*_*i*_) enter a structural trunk comprising six invariant point attention (IPA) blocks per pass, with residue and rigid-frame updates. First-pass binary pair probabilities are obtained through a learned pair head, detached and embedded into the pair pathway for a second pass with shared trunk parameters. At each flow evaluation, the second pass produces refined residue and pair states together with backbone-frame predictions for the sampler. **c**, Soft pair-biased sequence decoding. Binary-pair logits are projected into additive self-attention biases, while conditional pair-type probabilities are weighted by binary pairing probabilities before a separate bias projection. Four applications of a shared Transformer block produce per-residue nucleotide probabilities, from which the RNA sequence is decoded. Pair colours distinguish the predicted AU, GC, GU and OTHER classes.

We evaluated RiboCouple on generated and experimentally determined RNA backbones, assessing sequence–structure compatibility across multiple structure predictors, transfer to independently generated backbones, and pair-confidence-based candidate prioritization. We further tested whether the decoding gains depend on interaction information aligned to the recipient backbone.

## Results

### Interaction-conditioned decoding improves RNA backbone–sequence compatibility

We first asked whether transferring the inferred pair state improves sequence design for generated RNA backbones. The shared cohort comprised 550 sequence-free backbones produced by the RiboCouple flow sampler (50 at each length from 50 to 150 nucleotides in increments of 10). Three independently trained sequence decoders were compared: pair-type decoding used binary and typed pair-derived attention biases, binary-pair decoding used only binary pair bias, and topology-free decoding omitted pair-derived attention-bias projections (Methods). On these shared backbones, pair-type decoding achieved a joint-validity rate of 45.1%, compared with 41.3% for binary-pair decoding and 33.5% for topology-free decoding (Fig. 2a). The pair-type improvement over topology-free decoding was 11.6 percentage points (95% target-bootstrap CI, 8.4–15.1 percentage points). Generated sequences were independently refolded with RhoFold+ [8], and joint validity required TM-score at least 0.45 and root mean square deviation (RMSD) at most 5.5 Å.

**Figure 2.**
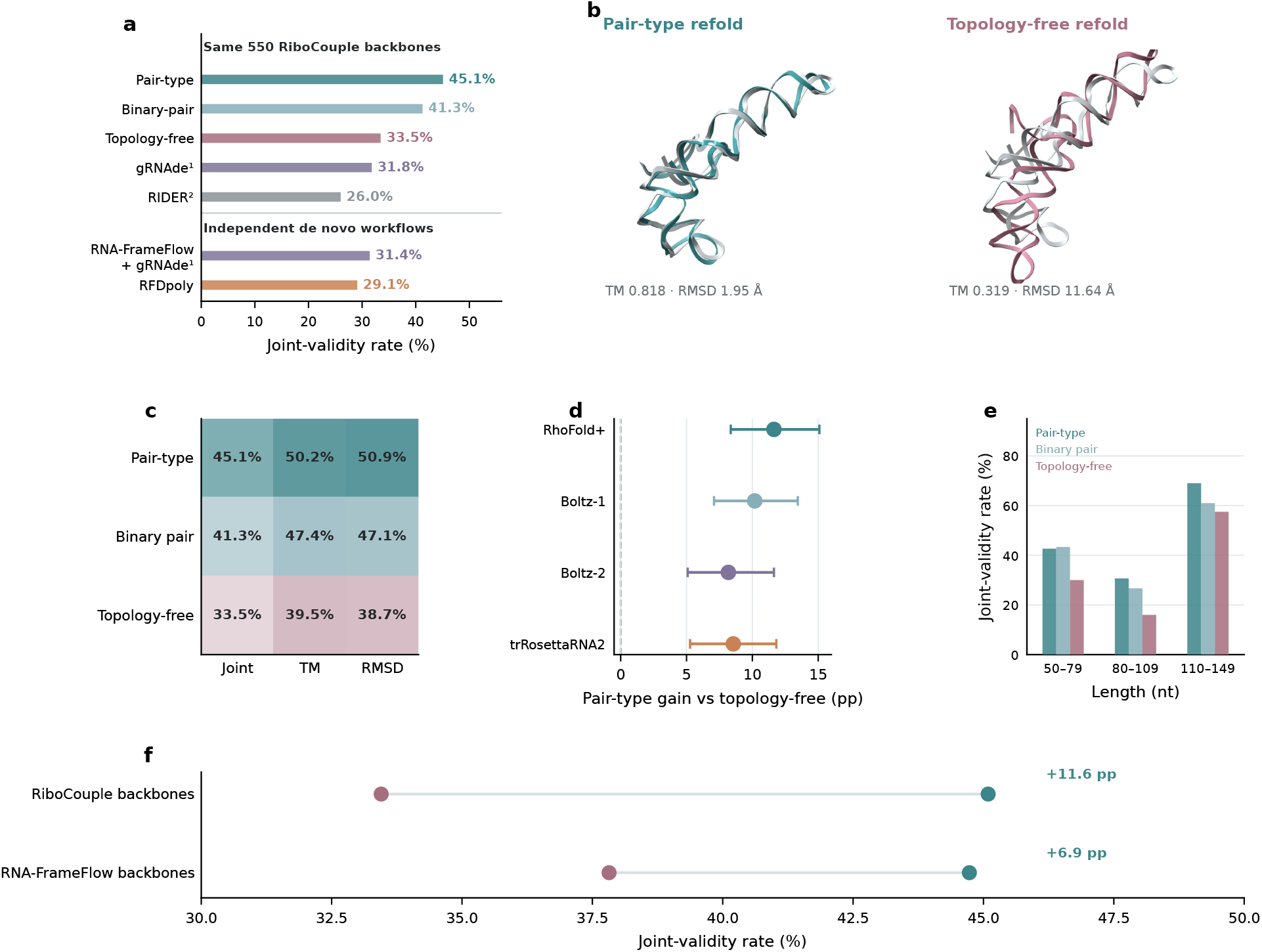
Pair-derived attention bias improves RNA backbone–sequence compatibility. Joint validity requires both TM-score *≥* 0.45 and RMSD *≤* 5.5 Å. **a**, Joint-validity rates for three RiboCouple decoders (pair-type, binary-pair and topology-free) and two external sequence-design methods (gRNAde^1^ and RIDER^2^) evaluated on the same 550 RiboCouple-generated backbones (upper group). The lower group shows length-matched independent de novo workflows (RNA-FrameFlow + gRNAde^1^ and RFDpoly). ^1^ Expected single-sample joint-validity rate averaged across eight fixed candidate positions. ^2^ One sequence output per backbone. **b**, Representative refolded structures for pair-type and topology-free decoding. **c**, Validity rates across joint, TM-score and RMSD criteria for the three RiboCouple decoder variants. **d**, Paired pair-type-minus-topology-free joint-validity differences across RhoFold+, Boltz-1, Boltz-2 and trRosettaRNA2, all evaluated on the same 550 backbones; intervals show 95% paired target-bootstrap confidence intervals. **e**, Length-stratified joint-validity rates for pair-type, binary-pair and topology-free decoding across 50–79, 80–109 and 110–149 nt. **f**, Absolute joint-validity rates for pair-type and topology-free decoding on RiboCouple and RNA-FrameFlow backbone cohorts; within each cohort, the line connects aggregate rates for the two decoders evaluated on the same backbone targets, and text labels report the pair-type increment in percentage points. Exact two-sided McNemar tests for the pair-type-minus-topology-free difference gave *P* = 1.15 *×* 10^*−*11^ on RiboCouple backbones and *P <* 10^*−*4^ on RNA-FrameFlow backbones.

The improvement persisted across all nine combinations of TM-score and RMSD thresholds tested, with pair-type gains of 10.2–12.2 percentage points (Supplementary Table S3b), and across RhoFold+, Boltz-1, Boltz-2 and trRosettaRNA2 [24], with gains of 8.18–11.64 percentage points (Fig. 2d and Supplementary Table S4). A representative pair-type versus topology-free structural comparison is shown in Fig. 2b.

We next applied the three RiboCouple decoders to a common cohort of 550 independently generated RNA-FrameFlow backbones. Joint validity was 44.7%, 44.2% and 37.8% for pair-type, binary-pair and topology-free decoding, respectively. The pair-type-minus-topology-free difference was 6.9 percentage points (95% paired target-bootstrap CI, 4.2–10.0 percentage points).

For comparison with an independent de novo design workflow, we evaluated RFDpoly [27] designs generated with the released RNA-only checkpoint on a length-matched set of 550 designs, using the same frozen RhoFold+ refolding, TMscore alignment and joint-validity criteria. RiboCouple pair-type decoding achieved mean TM-score 0.456 and joint validity 45.1%, compared with TM-score 0.308 and joint validity 29.1% for RFDpoly (Fig. 2a and Supplementary Table S5d). RFDpoly had higher joint validity at 70–100 nt, whereas RiboCouple had higher joint validity at 50, 60 and 110–140 nt.

For fixed-backbone sequence-design context on the same RiboCouple backbones, gRNAde achieved an expected single-sample joint-validity rate of 31.8% and RIDER a single-output rate of 26.0% (Fig. 2a and Supplementary Table S3). On RNA-FrameFlow backbones, the corresponding expected single-sample rate was 31.4%. Together, these results show that interaction-conditioned decoding improves computational sequence–structure compatibility on RiboCouple-generated backbones and on those produced independently by RNA-FrameFlow.

### Decoding depends on backbone alignment of the transferred relational state

We next asked whether decoding benefited from pair information in aggregate or from pair information aligned to the specific backbone. Across 110 recipient backbones, replacing the binary and typed pair logits supplied to the frozen decoder with those from four exact-length, global-statistics-matched donor backbones reduced joint validity from 46.4% to 21.4% (aligned-minus-donor difference, 25.0 percentage points; 95% nested-bootstrap CI, 16.4–33.6 percentage points) and mean TM-score (calculated with US-align) from 0.447 to 0.319 (Fig. 3d), while simple sequence-composition diagnostics remained similar. Thus, matching global pair-state statistics was not sufficient; decoding depended on whether the transferred pair state was aligned with the recipient backbone.

**Figure 3.**
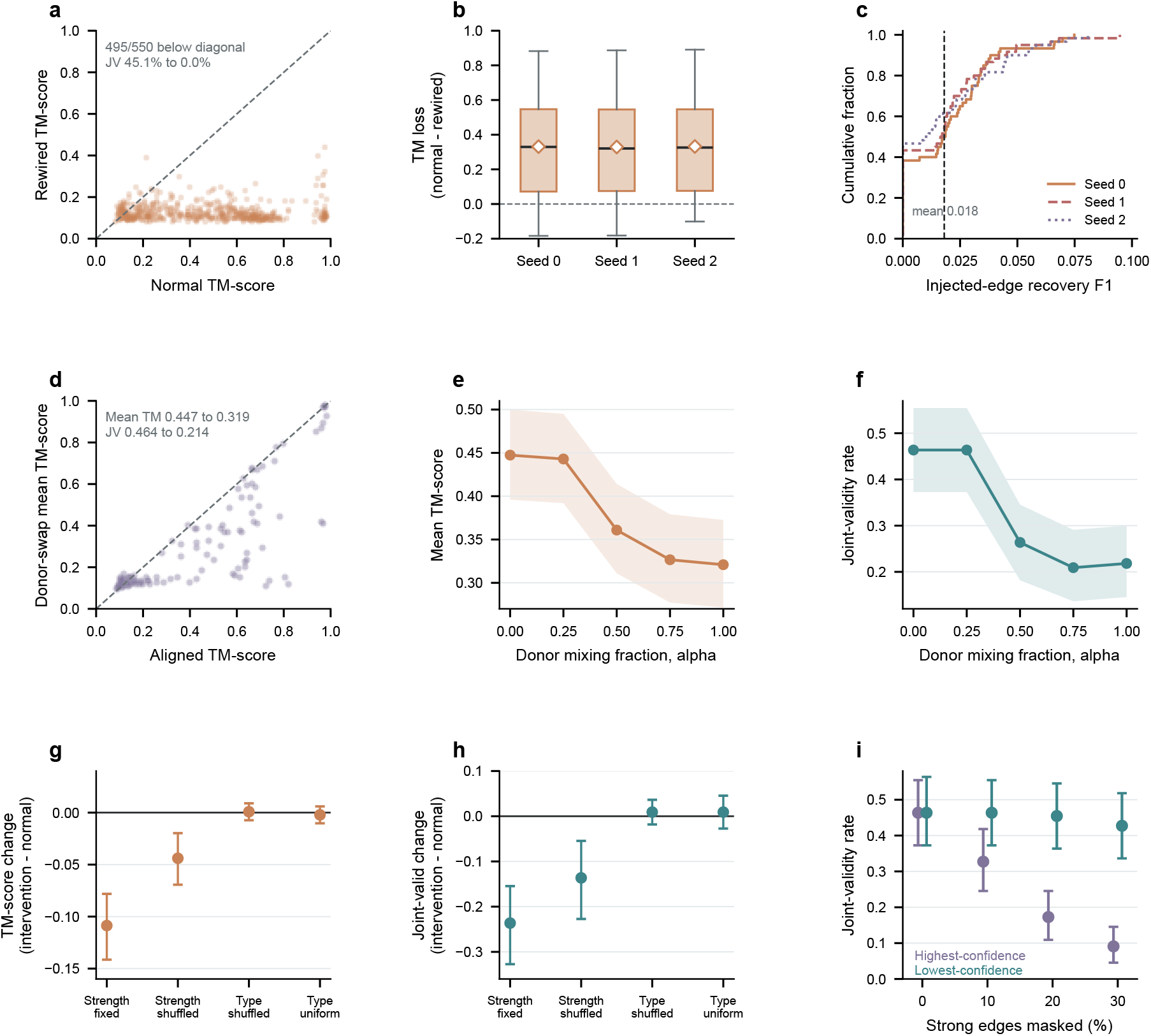
Structured interventions probe the contribution of backbone-matched pair information to sequence decoding. **a**, Normal versus rewired TM-scores on 550 shared backbones for one intervention seed. Across three seeds, 495–497 of 550 targets have lower TM-score after rewiring, and joint validity decreases from 45.1% to 0.0%. **b**, Per-backbone TM-score loss, defined as normal minus rewired, across three intervention seeds on the same 550 backbones. Boxes span the interquartile range, black lines show medians and white diamonds show means; whiskers extend to the most extreme observations within 1.5 interquartile ranges. **c**, Cumulative distributions of recovery F1 for deliberately rewired edges after refolding across three intervention seeds; mean injected-edge F1 was 0.018. **d**, Aligned TM-score versus the mean TM-score across four independently decoded exact-length, global-statistics-matched donor substitutions for each of 110 backbones; 80 targets fall below the identity line. Joint validity decreases from 46.4% to 21.4%. **e**,**f**, Mean TM-score (**e**) and joint-validity rate (**f** ) during interpolation from the aligned pair state toward a donor pair state; shaded bands show 95% target-bootstrap confidence intervals and lines connect measured interpolation levels. **g**,**h**, Paired changes in TM-score (**g**) and joint-validity rate (**h**) after fixing or shuffling pair strengths and shuffling or uniformizing conditional pair classes; joint-validity differences are in percentage points (intervention minus normal); error bars show 95% paired target-bootstrap confidence intervals. Fixing all binary pair logits at zero (pairing probability 0.5) caused the largest mean deterioration among the tested interventions. **i**, Joint-validity rate after masking increasing fractions of the highest- or lowest-confidence strong edges; horizontally offset point–interval marks show 95% target-bootstrap confidence intervals. Dashed diagonals in **a**,**d** denote equality. TM-scores were computed with TMscore in **a**,**b** and US-align in **d–i.** Joint validity requires TM-score *≥* 0.45 and RMSD *≤* 5.5 Å, with RMSD for **d–i** taken from the corresponding frozen TMscore evaluation (Methods).

A complementary rewiring experiment tested whether preserving coarse graph statistics was sufficient. On 550 backbones, reassigning residue partners by degree-preserving double-edge swaps with separation-aware matching and subsequent quality control reduced joint validity from 45.1% to 0.0% across all three intervention seeds; the deliberately rewired edges were rarely recovered after refolding (mean injected-edge recovery F1 = 0.018; Fig. 3a–c). Rewiring changes partner placement together with the background pair-logit input, since non-edge entries are set to zero logit.

Gradually mixing donor pair logits into the backbone-aligned state reduced structural recovery. Mean TM-score (calculated with US-align) declined with increasing donor contribution, while joint validity dropped sharply when the donor fraction reached 0.50 (Fig. 3e,f).

Component-level perturbations further identified the pair information used by the decoder. Among the tested fixed-decoder interventions, setting all binary pair logits to zero (corresponding to a pairing probability of 50%) caused the largest deterioration in TM-score and joint validity; shuffling binary pair strengths was also detrimental, whereas shuffling or uniformizing the conditional pair-type channel produced little average change (Fig. 3g,h). Masking the highest-confidence strong edges was substantially more damaging than masking an equal fraction of the lowest-confidence edges (Fig. 3i). These findings identify a strong inference-time dependence on correctly placed, high-confidence pair relations.

### Recycling refines soft pair guidance used in sequence decoding

To assess whether recycling improved the pair representation itself, we compared first- and second-pass pair probabilities on the 94-chain native benchmark. Recycling increased the area under the precision–recall curve (AUPRC) from 0.649 to 0.673 and improved pair-probability calibration, although precision at 80% recall declined (Fig. 4a, Extended Data Fig. 4 and Supplementary Table S8b).

**Figure 4.**
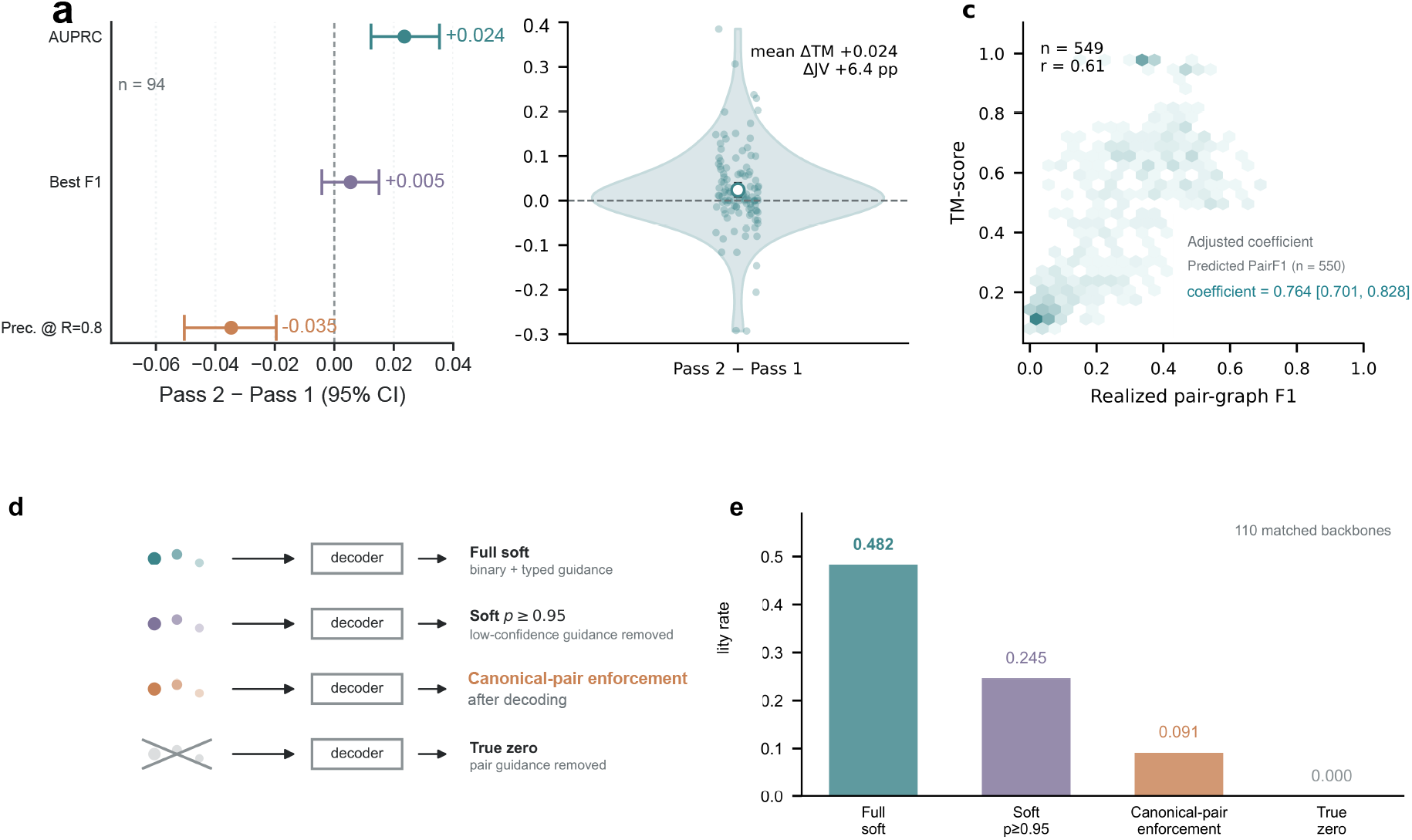
Pair recycling refines pair guidance used during sequence decoding. **a**, Pass-2-minus-Pass-1 changes in AUPRC, best F1 and precision at 80% recall on the 94-chain native benchmark; intervals are 95% target-bootstrap CIs. **b**, Target-level TM-score change calculated with US-align when the same decoder receives second-pass rather than first-pass binary pair-logit inputs while all remaining decoder inputs are fixed; the mean change is +0.024, joint validity increases by 6.4 percentage points, and the open circle marks the mean. **c**, Association between realized pair-graph F1 and downstream TM-score calculated with US-align. Realized F1 compares the thresholded (*p >* 0.5), symmetrized pair graph with hydrogen-bonded pairs in the refolded structure (*n* = 549; one design had no edge above threshold). The annotated adjusted coefficient uses PairF1 recomputed in a separate forward pass on all 550 targets, controlling for RNA length, pair density, backbone novelty and reference-sequence refoldability. **d**, Four decoding strategies in a separately sampled 110-backbone cohort: full-soft attention guidance, confidence-gated soft guidance retaining edges with *p ≥* 0.95, deterministic canonical-pair enforcement after decoding and decoding without pair-derived attention biases. **e**, Corresponding joint-validity rates of 48.2%, 24.5%, 9.1% and 0.0%, respectively.

To test how recycling affects downstream sequence decoding, the same decoder was supplied with either first-pass or second-pass binary pair logits while the backbone and all remaining decoder inputs were held fixed. Second-pass logits improved mean TM-score by 0.024 (calculated with US-align) and increased joint validity by 6.4 percentage points (Fig. 4b and Extended Data Fig. 2). Across 549 evaluable designs (one of the 550 designs had no predicted pair edge above the *p >* 0.5 threshold), realized pair-graph F1 was positively associated with downstream TM-score (*r* = 0.607; Fig. 4c). A separate adjusted analysis using PairF1 recomputed in another forward pass on all 550 targets found a positive association after accounting for RNA length, pair density, backbone novelty and reference-sequence refoldability (Supplementary Table S6b).

Four pair-guidance conditions were compared in a separately sampled 110-backbone cohort using the same frozen pair-trained decoder: full-soft attention guidance, confidence-gated soft guidance, post-decoding deterministic canonical-pair enforcement and complete removal of pair-derived guidance. Joint validity was 48.2% with full-soft guidance, 24.5% with confidence-gated soft guidance retaining only edges with *p ≥* 0.95, 9.1% with post-decoding deterministic canonical-pair enforcement and 0.0% with pair guidance removed (Fig. 4d,e). Confidence-gated soft guidance retained intermediate performance, whereas post-decoding deterministic enforcement performed substantially worse than full-soft guidance.

### Pair-coupled decoding improves secondary-structure agreement on native backbones

We evaluated pair-coupled decoding on 94 experimentally determined RNA backbones from a temporally held-out evaluation cohort, using inferred residue partners to coordinate nucleotide choices under the fixed model (the reciprocal-best non-overlapping pair-selection and joint-assignment procedure is detailed in the Supplementary Methods). Compared with residue-wise argmax on the identical 94 chains, pair coupling changed full-length sequence recovery from 43.52% to 41.57%, secondary-structure Matthews correlation coefficient (MCC) from 0.2660 to 0.5462 and canonical base-pair F1 (BP-F1) from 0.2672 to 0.5414 (Fig. 5a–c and Supplementary Table S9b). BP-F1 used the same X3DNA-DSSR-derived canonical-pair reference for every method. Pair coupling therefore traded a small decrease in sequence recovery for markedly improved secondary-structure agreement. The within-target paired gain was 0.280 for MCC (95% target-bootstrap CI, 0.225–0.338) and 0.274 for BP-F1 (95% CI, 0.220–0.328); per-chain win counts and two-sided Wilcoxon tests for all three metrics are reported in Supplementary Table S9b. The MCC improvement was also consistent under PDB-entry-level resampling (Supplementary Table S9b). Alternative pair-constraint decoding strategies, including binarized predicted and native-reference pair graphs, are compared on the same 94 chains in Supplementary Table S9c.

**Figure 5.**
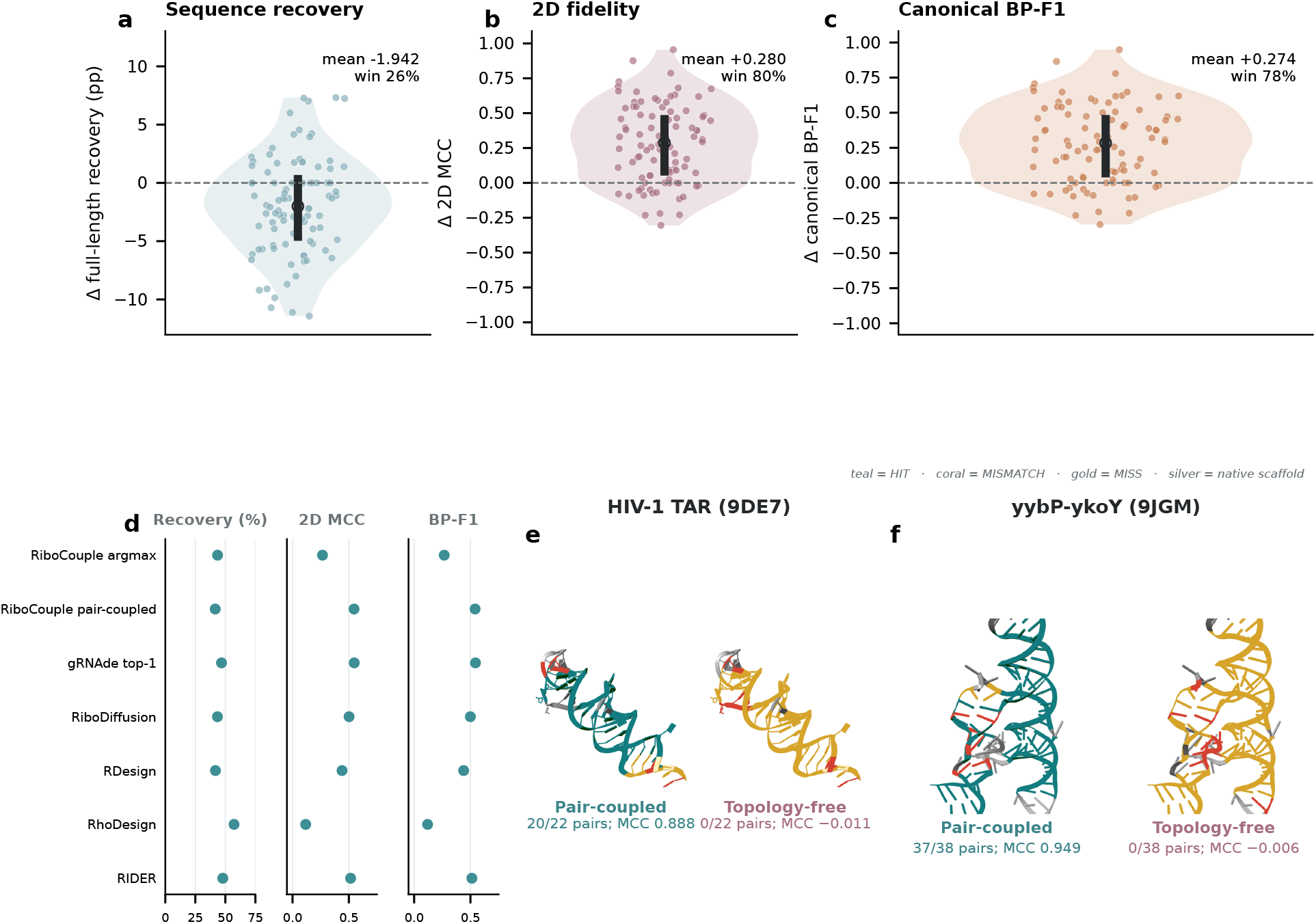
Pair-coupled decoding improves secondary-structure agreement on native backbones. **a–c**, Distributions of pair-coupled minus residue-wise-argmax changes on the same 94 native RNA backbones in full-length sequence recovery (**a**), 2D MCC (**b**) and canonical BP-F1 (**c**). Effect estimates, confidence intervals, win counts and statistical tests are reported in Supplementary Table S9b. **d**, Fixed-backbone single-output benchmark of RiboCouple argmax and pair-coupled decoding, gRNAde top-1, RiboDiffusion, RDesign, RhoDesign and RIDER, separately comparing recovery, 2D MCC and canonical BP-F1 on a common denominator of 94 targets. **e**,**f**, Illustrative native-structure pair-realization examples for HIV-1 TAR (9DE7) and the yybP-ykoY riboswitch (9JGM).

External inverse-folding methods exhibited different trade-offs on the same 94-chain benchmark (Fig. 5d and Supplementary Table S9). At one sequence per target, RiboCouple pair-coupled decoding yielded MCC 0.5462 and canonical BP-F1 0.5414, close to gRNAde top-1 (0.5479 and 0.5448), with higher secondary-structure agreement than RIDER (0.5158 and 0.5127), RiboDiffusion (0.5021 and 0.4995), RDesign (0.4398 and 0.4397) and RhoDesign (0.1166 and 0.1194). RhoDesign achieved the highest sequence recovery (57.23%).

Among 65 of the 94 benchmark chains whose nearest RiboCouple training-cluster representative had a US-align TM similarity of at most 0.45, pair-coupled decoding increased MCC by 0.259 (95% paired target-bootstrap CI, 0.195–0.323) and BP-F1 by 0.252 (95% CI, 0.190–0.314). Across all 94 chains, the gain was positively associated with nearest-training-cluster similarity (Spearman *ρ* = 0.277, *P* = 0.007 for MCC; *ρ* = 0.287, *P* = 0.005 for BP-F1).

HIV-1 TAR (9DE7) and the yybP-ykoY riboswitch (9JGM) illustrate pair realization in separate native-structure case studies (Fig. 5e,f): in the predicted secondary structures of the designed sequences, pair-coupled decoding reconstructed 20 of the 22 native canonical pairs of TAR, including the upper-stem pairs adjoining the UCU bulge that mediates Tat recognition [28, 29], and 37 of the 38 native pairs of the yybP-ykoY aptamer, whose pairing network scaffolds the Mn^2+^-sensing conformation required for riboswitch switching [30, 31].

Across the 94 native chains, the mean per-chain canonical compatibility at sequence-distant native pairs (sequence-index separation |*i − j*| *≥* 10; 1,646 pairs in total) increased from 74.7% with residue-wise argmax to 96.3% with pair-coupled decoding. Positive pooled gains were observed across all 12 annotated RNA groups, including ribozymes and IRES elements, and canonical compatibility reached 99.4% for tRNA pairs (Supplementary Tables S9d and S9e).

### Pair confidence prioritizes backbones before external refolding

We derived a pre-refolding confidence score from the pair representation by summing the ten largest binary pair probabilities over directed off-diagonal pairs (*i < j*). Across four score strata formed without splitting tied values, joint validity increased from 14.7% in the lowest stratum to 64.7% in the highest stratum (Fig. 6a,b). Under tie-aware selection of the *K* = 248 highest-ranked candidates by pair confidence (with randomized resolution of cutoff ties), pair-type joint validity was 59.6%, compared with 46.7% for topology-free decoding on the same selected backbones; the pair-type-minus-topology-free difference remained 12.8–13.6 percentage points across *K* = 50–248 (Fig. 6c). The same score enriched pair-type outcomes across all four structure evaluators (Fig. 6d). Applied before refolding to independently generated backbones, it increased joint validity from 44.7% to 62.0% on RNA-FrameFlow backbones and from 21.8% to 37.1% on RFDpoly backbones at *K* = 248 after RiboCouple pair-type decoding (Fig. 6e). These results identify pair confidence as a pre-refolding ranking signal, while the persistent pair-type advantage within selected sets indicates a separate contribution of pair-conditioned decoding.

**Figure 6.**
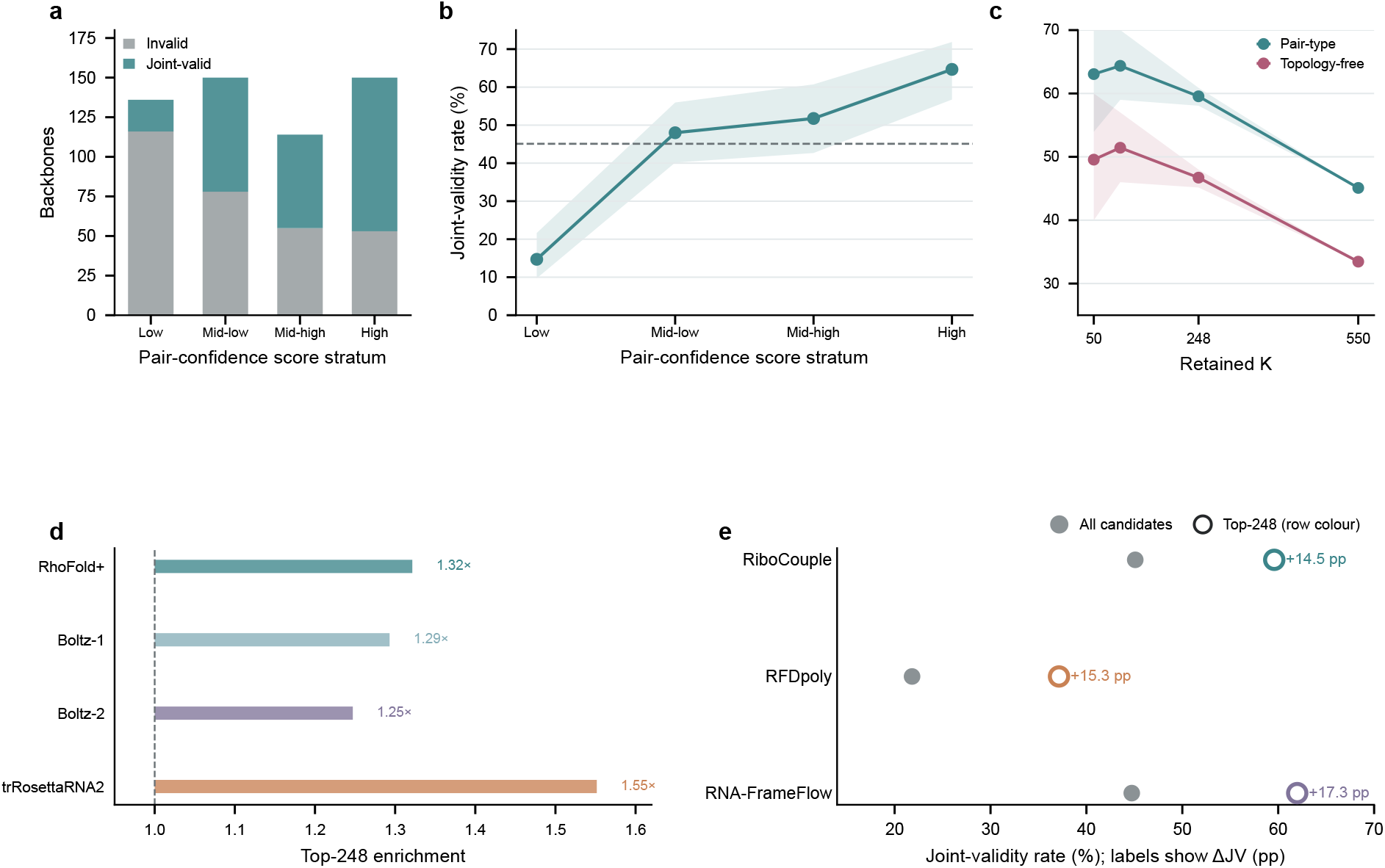
Pair-head confidence prioritizes backbones before external refolding. The confidence score is the pre-refolding sum of the ten largest binary pair probabilities over valid off-diagonal residue pairs, taking the directed entry with *i < j* for each unordered pair. **a**, Numbers of invalid and joint-valid RiboCouple-generated backbones across four low-to-high score strata, formed by grouping complete score-tie groups without splitting equal values (from low to high, *n* = 136, 150, 114 and 150). **b**, Joint-validity rates in the same strata; bands are 95% Wilson confidence intervals. **c**, Pair-type and topology-free joint-validity rates across retained-backbone counts on the same 550 RiboCouple backbones; bands show 95% intervals from tie-aware selection (at *K* = 248, 58.1–60.9% for pair-type and 45.2–48.0% for topology-free). **d**, Top-248 joint-validity enrichment across four independent structure evaluators. **e**, Joint-validity rates for all candidates (grey) and tie-aware top-248 selection (unfilled colored circles) after the RiboCouple pair-type decoder was applied to RiboCouple, RFDpoly and RNA-FrameFlow backbones; labels report absolute ΔJV in percentage points.

## Discussion

RiboCouple establishes a probabilistic, backbone-derived interaction topology that transfers backbone-aligned relational information into nucleotide decoding through soft attention biases. Across generated and experimentally determined backbones, pair guidance improved computational sequence–structure compatibility and supplied a pre-refolding signal for prioritizing backbones likely to satisfy the computational validity criteria.

The interventions provide different evidence for backbone alignment and typed conditioning. Donor substitution, partner rewiring and confidence-ranked masking showed that decoding depends on correctly placed, high-confidence pair relations. Acute perturbation of conditional pair types produced little average change, whereas independently trained pair-type decoding achieved higher joint validity than binary-pair decoding (45.1% versus 41.3%) and topology-free decoding (33.5%). This interface retained utility beyond RiboCouple’s own backbone generator. Pair-conditioned decoding improved joint validity on backbones from an independent generator (RNA-FrameFlow), and pair-coupled decoding improved secondary-structure agreement on experimentally determined backbones. Pair-head confidence enriched computationally valid candidates on RiboCouple-generated backbones and, after frozen RiboCouple decoding, on independently generated RNA-FrameFlow and RFDpoly backbones. The ranking signal was also observed across independent structure evaluators. Because pair confidence can be computed before any external structure prediction, pre-ranking offers a way to reduce computational screening effort: at the observed top-248 enrichments (1.25*×*– 1.55*×* across four evaluators), this corresponds to approximately 20–35% fewer external refolding evaluations per computationally valid design, excluding the cost of ranking (Supplementary Table S9g).

Together, these findings show that backbone-derived pair information can guide RNA sequence decoding and inform candidate selection before external refolding.

## Methods

### RiboCouple model overview

RiboCouple comprises a backbone-generation stage and an interaction-conditioned sequence-decoding stage. During each flow-model evaluation, an SE(3)-equivariant trunk processes the current residue frames while maintaining residue and pair representations. Binary pair probabilities predicted in the first pass are recycled through the shared trunk in a second pass to refine geometric and relational states. The refined pair representation then produces binary-interaction and conditional pair-type signals that are converted to additive attention biases for a Transformer sequence decoder. The sequence-decoding model is initialized from the pair-recycled structural checkpoint; during the final decoding stage, the structural trunk and binary interaction head are held fixed while the pair-type head and Transformer decoder are optimized.

### Study design

RiboCouple was evaluated in two complementary settings. The generated-backbone benchmark tested whether sequences decoded for newly generated RNA backbones refolded into compatible three-dimensional structures. The native-backbone benchmark tested sequence recovery and canonical secondary-structure compatibility on experimentally determined RNA structures. Pair-type, binary-pair and topology-free variants were evaluated using shared backbones and matched inference settings. Unless stated otherwise, controlled inference-time pair-state interventions held the checkpoint, backbone coordinates, sequence decoder and decoding seeds fixed and changed only the pair-derived signals supplied at inference time.

### Model architecture and training

#### RNA structural data and backbone representation

Training structures were drawn from the RNA-Solo snapshot released on 31 October 2023, a frozen PDB-derived RNA dataset [32, 33], and restricted to single RNA chains of 40–150 nucleotides resolved at 4.0 Å or better, yielding 5,038 training examples. Following the RNA-FrameFlow parameterization [11], each nucleotide was represented by a rigid frame *T*_*i*_ = (*R*_*i*_, *x*_*i*_) ∈ *SE*(3): *x*_*i*_ places C4^*′*^ in global coordinates and *R*_*i*_ orients the local frame defined by the C3^*′*^–C4^*′*^–O4^*′*^ sugar triad. C3^*′*^, C4^*′*^ and O4^*′*^ are ribose atoms whose non-collinear positions define the local coordinate system. Eight torsion angles specify rotations outside this frame. The model therefore does not predict all PDB coordinates independently. For atom *a* in residue *i*, the reconstructed coordinate has the form

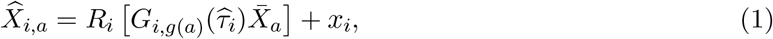

where 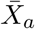 is an atom coordinate in the fixed local nucleic-acid template and *G*_*i,g*(*a*)_ is the torsion-dependent transform for the atom’s rigid group. Dataset composition, frame construction and the complete reconstruction procedure are given in the Supplementary Methods.

#### DSSR-derived base-pair annotations

Base-pair supervision was prepared separately for this study. Experimental RNA structures in the training set were analysed with DSSR [34], and the resulting residue-pair annotations were aligned to the residue indices used by the model. For each structure, we stored a binary base-pair matrix together with annotation metadata and a label mask indicating whether the DSSR output could be aligned reliably. These annotations were generated before model training and read directly by the data loader.

Pair-type labels were constructed online from the native nucleotide identities and the DSSR-derived binary matrix. Positive pairs were assigned to AU, GC, GU or OTHER: AU and GC are canonical Watson–Crick classes, GU is the common wobble class, and OTHER collects the remaining DSSR-annotated nucleotide-pair compositions. Valid residue-index pairs without DSSR-annotated interactions served as negative examples for the binary-pair objective but were excluded from the conditional pair-type objective; invalid residues and samples without reliable pair annotations were also excluded from the pair-type objective. Thus, the binary task identifies base-pair partners, whereas the typed task predicts the nucleotide-pair class at annotated positive edges. During training, pair-type supervision was applied only at annotated positive pair positions. During inference, the predicted binary-pair probability and conditional pair-type distribution were retained as continuous quantities and supplied to the sequence decoder as attention biases rather than as hard nucleotide constraints.

#### SE(3)-equivariant flow-matching backbone model

Following flow-matching formulations [11, 35, 36], an RNA of length *L* is represented by a continuous structural state comprising residue frames **T**_*t*_ *∈ SE*(3)^*L*^. Initial reference translations and rotations are connected to data frames by a linear path in ℝ^3^ and a geodesic path on *SO*(3), respectively:

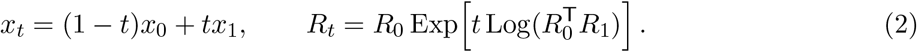

The invariant-point-attention (IPA) trunk [37] receives **T**_*t*_, flow time, residue states and pair states and predicts the terminal translations and rotations that parameterize the translational and rotational vector fields. A torsion head predicts the eight internal rotations used for atom reconstruction; torsions are structural outputs but are not themselves integrated as part of the *SE*(3) flow state. Auxiliary objectives act on reconstructed backbone atoms, inter-residue distances and torsion vectors. This continuous geometry and the associated residue and pair representations provide the inputs to the interaction and sequence modules.

#### Binary interaction prediction

A binary pair head maps the trunk pair representation for residues *i* and *j* to a logit 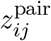. The binary objective was evaluated over valid, non-diagonal residue pairs,

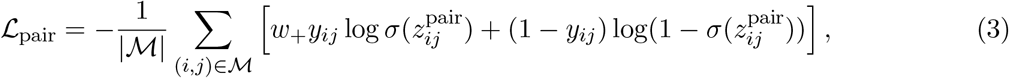

where *y*_*ij*_ is the DSSR-derived binary label and *ℳ* is the valid pair mask. Class balancing was used to account for the large excess of negative residue pairs.

#### Pair-type prediction

The pair-type head produces four logits 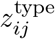 for AU, GC, GU and OTHER. Cross-entropy was evaluated only on DSSR-annotated positive pairs,

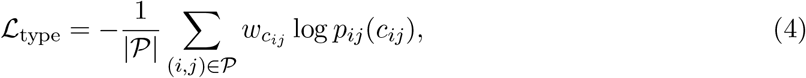

where *P* is the set of valid positive pairs and *c*_*ij*_ is the corresponding pair class. Class frequencies were used to balance the pair-type objective across the classes represented in each batch.

#### Pair recycling

At each structural evaluation, RiboCouple applies the shared structural trunk twice. The first pass produces residue and pair representations together with binary pair logits. The corresponding probabilities are detached, embedded and fed back into the pair pathway for a second pass. This second pass updates the residue features, pair features and rigid states; its outputs define the final structure, the reported pair predictions and the pair-derived attention biases supplied to the sequence decoder. The binary-pair objective was evaluated on the second-pass logits, and no thresholding was applied between the two passes. For fixed-backbone sequence decoding, supplied backbone coordinates are processed by this same full two-pass forward computation; no diffusion-trajectory state is consumed by the sequence decoder.

#### Pair-biased Transformer sequence decoder

The sequence decoder applies a Transformer block [38] four times with shared parameters, using hidden dimension 256, eight attention heads and a dropout rate of 0.1. It receives 256-dimensional residue representations and additive pair-derived attention biases from the 128-dimensional pair representation. For attention head *h*, the unnormalized attention score between residues *i* and *j* is

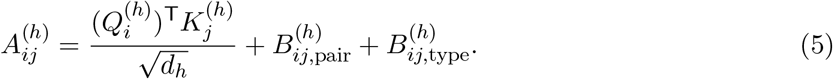

The binary pairing probability 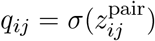 is combined with the conditional pair-type distribution **t**_*ij*_ = softmax 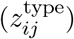 to form

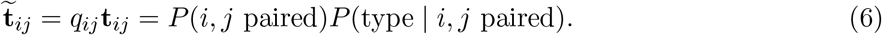

The additive bias terms in the attention score are computed by two separately learned, head-specific linear projections:

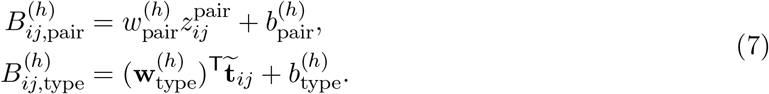

Here the scalar binary logit and four-dimensional soft typed feature enter different bias projections; the projections produce one additive score per attention head. Pair-type classes encode unordered nucleotide composition because the type channel represents pair composition, whereas residue-specific representations preserve positional identity and allow separate nucleotide logits for residues *i* and *j*. The Transformer outputs a categorical distribution over A, C, G and U at each valid residue, using the full soft binary and typed distributions throughout decoding. Three independently trained Transformer decoding conditions were used in the primary comparison: the pair-type decoder received both binary and typed pair-derived biases, the binary-pair decoder received only the binary pair bias, and the topology-free decoder contained no pair-derived attention-bias projection.

#### Training objective and generation

Training was performed in two stages. In stage 1, the backbone-generation model and binary interaction head were optimized with

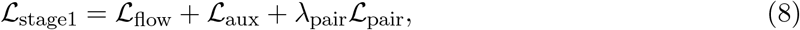

where *L*_flow_ comprises translational and rotational terms weighted 2.0 and 1.0, respectively, *L*_aux_ has weight 1.0, and *λ*_pair_ = 1.0. This stage produced the pair-recycled structural checkpoint.

In stage 2, each sequence-decoding condition was initialized from this checkpoint. The structural trunk and binary interaction head were frozen, and only the pair-type head, when present, and the Transformer sequence decoder were optimized. The stage-2 objective was

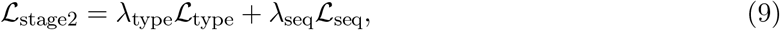

where *L*_seq_ is nucleotide cross-entropy over valid residues. We used *λ*_type_ = 0.2 and *λ*_seq_ = 1.0; for the binary-pair and topology-free conditions, the pair-type term was omitted. The pair-type, binary-pair and topology-free sequence heads were trained independently under their respective attention-bias configurations. Training used AdamW with a learning rate of 10^*−*4^, weight decay 0.01 and no learning-rate scheduler. All models were trained on a single node equipped with four NVIDIA GeForce RTX 4090 GPUs (24 GB memory each) using distributed data parallel (DDP) training, with a per-device batch size of 28 and at most 200 epochs. Candidate checkpoints for each sequence-decoding condition were selected using TM-validity on a separate unconditional-evaluation cohort, distinct from the frozen 550-backbone primary benchmark, with approximately ten generated backbones per evaluated length. The complete training and inference configuration is summarized in Supplementary Table S10b.

At inference, translations were initialized from a chain-centred Gaussian reference distribution and rotations from the uniform distribution on *SO*(3). The learned vector field transported these frames over 50 sampling steps beginning at *t* = 0.01. Rotation used a linear training schedule and an exponential sampling schedule with rate 10; translation used linear schedules for training and sampling. The final frames and predicted torsions were converted to atomic coordinates by the reconstruction above. In parallel, first-pass binary interaction probabilities were detached and recycled through the second structural pass without thresholding; the final binary-interaction and pair-type logits then supplied two soft, pair-derived attention biases to the Transformer, which generated one A/C/G/U distribution per valid residue.

### Evaluation benchmarks and metrics

#### Generated-backbone compatibility benchmark

The primary benchmark comprised 550 RiboCouple-generated backbones shared across the pair-type, binary-pair and topology-free decoders. Each decoder produced one residue-wise argmax sequence per backbone for this benchmark. Each sequence was independently refolded with RhoFold+ [8], and the predicted structure was compared with its design backbone using TM-score and RMSD computed by TMscore [39], following the RNA-FrameFlow self-consistency protocol [11]. TM-validity denotes the fraction of designs with TM-score at least 0.45, RMSD-validity denotes the fraction with RMSD at most 5.5 Å, and joint validity requires both criteria. Matched-adversarial rewiring used the same TMscore-derived TM-score and RMSD fields as the primary benchmark. For the representation-intervention and recycling analyses, TM-score was calculated with US-align, whereas RMSD was taken from the corresponding frozen TMscore evaluation field; joint validity always required both thresholds. Details of the frozen-cohort protocols are given in Supplementary Methods. External inverse-folding controls were also evaluated on this same 550-backbone RiboCouple cohort. For gRNAde, eight independently sampled candidate sequences were evaluated per backbone; the reported single-sample statistic is the mean over the eight fixed candidate positions, estimating the expected outcome of one draw rather than selecting the best-of-eight candidate. RIDER was evaluated in single-output mode and scored with the same downstream refolding and validity definitions.

#### Cross-generator decoder evaluation

To test whether the learned sequence decoder remained usable outside the RiboCouple-generated backbone distribution, the frozen pair-type, binary-pair and topology-free decoders were applied to the same 550 RNA-FrameFlow backbones. Decoder checkpoints and backbone files were held fixed across conditions. Joint-validity contrasts were computed on matched targets.

#### Independent structure evaluators and length analysis

On the primary 550 RiboCouple-generated backbone cohort, pair-type, binary-pair and topology-free sequences were additionally evaluated with Boltz-1 [40], Boltz-2 [41] and trRosettaRNA2 [24] using the same target pairing. RhoFold+ was the frozen primary evaluator. Length-stratified RhoFold+ results were reported for 50–79, 80–109, 110–149 and 150 nucleotides.

#### Native-backbone sequence-design benchmark

Transfer to experimentally determined structures was evaluated on a common 94-chain temporally held-out native-backbone cohort. The first public release date of each of the 94 chains (earliest, 8 November 2023) postdated the training-data cutoffs of the compared model checkpoints. Sequence recovery was defined as per-residue identity over the full native target length. Each compared method contributed one sequence per target: RiboCouple argmax and pair-coupled decoding, gRNAde top-1, RiboDiffusion, RDesign, RhoDesign and RIDER. For gRNAde, top-1 denotes the first sampled candidate of its 16-sample inference protocol (a fixed sample index). All reported methods and metrics share the same *n* = 94 denominator.

Canonical base-pair F1 was measured under a common RNAfold protocol [42] by comparing canonical reference base pairs obtained from X3DNA-DSSR with canonical pairs in the ViennaRNA minimum-free-energy secondary structure predicted for each designed sequence. The 2D MCC was calculated over all unordered residue pairs using the same reference and predicted pair sets. The per-target metric definitions and calculation scripts are provided in the evaluation manifest.

Pair-coupled decoding was evaluated separately on native backbones as a fixed-model inference intervention, using reciprocal-best non-overlapping pairs selected from symmetrized pair probabilities; the complete selection and joint-assignment rule is provided in the Supplementary Methods and source data.

#### Predicted-to-realized pair-graph agreement

Hydrogen-bonded residue pairs were extracted from refolded structures and compared with the thresholded pair graph supplied to the sequence decoder. Precision, recall and F1 were calculated for each design from residue-pair identities and then summarized across targets. The thresholded realization scatter used 549 evaluable designs because one backbone had no edge above the pair-head threshold. Chain-length-adjusted and length-stratified analyses tested whether the raw association persisted after accounting for RNA length.

A complementary multivariable linear regression assessed the association between PairF1 and TM-score after adjustment for RNA length, pair density, backbone novelty and reference-sequence refoldability. Coefficients were estimated at the target level with nonparametric bootstrap confidence intervals. This adjusted analysis used 550 targets, whereas the thresholded realization scatter used 549 designs (Supplementary Methods and Supplementary Table S6b).

### Controlled interventions on interaction topology

#### Pair-guidance and deterministic-constraint interventions

We compared four ways of using the predicted relational state in a separately sampled, matched 110-backbone cohort while holding model parameters, backbone coordinates and decoding seeds fixed. The full-soft condition used binary and typed pair-derived attention guidance. In the confidence-gated soft condition, pair-derived guidance was retained only for edges with pair probability at least 0.95, with below-threshold binary and typed contributions removed from the attention pathway. In the true-zero condition, both binary and typed pair-derived attention signals were removed. For deterministic canonical-pair enforcement, the full-soft decoder first produced a sequence, after which an enforcement step derived from the predicted pair graph was applied.

For the matched-adversarial intervention, rewiring was constructed by degree-preserving double-edge swaps with separation-aware matching and subsequent quality control. The swaps changed interacting partners while targeting preservation of residue degrees, total pair count and the distribution of sequence separation |*i − j*|.

For injection, each rewired graph was converted to a pair-logit matrix by writing the model’s original predicted logit strengths onto the reassigned edge positions and setting all other entries to zero logit.

#### Matched donor relational-state substitution

Matched donor substitution was evaluated on the frozen 110-backbone intervention cohort. For each recipient backbone *A*, candidate donors were restricted to backbones of exactly the same length, and the four nearest donors were selected using

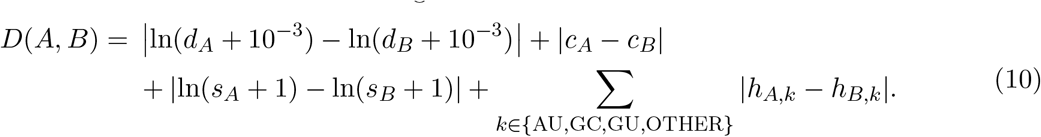

Here *d* = *n*_strong_*/L*^2^ is the density of residue-pair entries with sigmoid pair probability greater than 0.5, *c* is the mean sigmoid pair probability over these strong entries, *s* is their mean sequence separation |*i − j*|, and *h* is the normalized histogram of the argmax conditional pair-type classes over strong entries. The four terms were summed without additional weighting or standardization.

For each donor *B*, the recipient coordinates and second-pass residue representations from *A* were retained, whereas *B*’s second-pass raw binary pair logits and four-class conditional pair-type logits were substituted at the input of the frozen sequence head. The binary pair logits therefore entered the learned pair-bias projection directly, while the typed feature was formed inside the decoder as *σ*(*z*^pair^) softmax(*z*^type^) before its separate projection. The internal 128-dimensional pair representation was not exchanged. Each of the four donor states was decoded and refolded independently; donor outcomes were averaged only after evaluation. Sequence-composition diagnostics comprised GC fraction (the fraction of decoded nucleotides that were G or C), nucleotide-composition Shannon entropy and maximum homopolymer length. Uncertainty for the aligned-minus-donor joint-validity difference used a nested bootstrap that resampled recipient targets and donor outcomes.

#### Aligned-to-donor pair-state interpolation

For matched recipient–donor pairs, interpolation was performed on the raw binary-pair and conditional pair-type logits supplied to the frozen decoder,

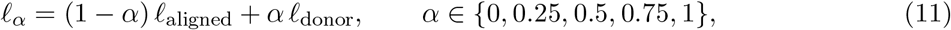

where *l* denotes either the binary pair logit or the four-component pair-type logit vector. Sequences were decoded and refolded independently at each interpolation level while the backbone, residue representations and decoder parameters were fixed.

#### Pair-strength and pair-type interventions

Component-level interventions used the same frozen 110-backbone cohort and sequence decoder. In the strength-fixed condition, the binary pair logits were set to zero, corresponding to sigmoid pair probability 0.5, while the conditional pair-type input was retained. In the strength-shuffled condition, binary pair-strength records were permuted across residue-pair entries while the conditional type input was retained. In the type-shuffled condition, conditional four-class pair-type records were permuted while the binary pair logits were retained. In the type-uniform condition, the conditional pair-type distribution was replaced by a uniform four-class distribution while the binary pair logits were retained. These interventions measured the frozen decoder’s sensitivity to changes in pair inputs.

#### Confidence-ranked edge masking

Within each target, strong pair relations were defined by sigmoid pair probability greater than 0.5 and ranked by binary pair confidence. The top or bottom 10%, 20% and 30% of these relations were masked at the sequence-head pair-state input while the backbone, frozen decoder and remaining relational information were held fixed. The matched masking fractions therefore compared decoder sensitivity to high- and low-confidence relations under the same intervention budget.

#### Evaluation of pair recycling

First- and second-pass pair representations were compared using pair-ranking AUPRC, best F1, precision at specified recall levels and probability-calibration metrics on matched native RNA targets. The contribution of recycled pair logits to sequence decoding was evaluated by replacing the second-pass binary pair-logit input with its first-pass counterpart while keeping the backbone, remaining decoder inputs and sequence decoder fixed, followed by paired comparison of joint validity and TM-score.

### Candidate ranking and structural diagnostics

#### Pair-based candidate prioritization

Pair-based confidence scores were matched to structural outcomes using stable target identifiers. For each backbone, the score was the sum of the ten largest binary pair probabilities over valid off-diagonal residue pairs, taking the directed entry with *i < j* for each unordered pair. Spearman correlation quantified its association with downstream TM-score using average ranks for tied scores. For each retained-set size *K*, candidates with scores strictly above the cutoff were retained and any remaining positions were sampled uniformly from targets tied at the cutoff. Tie-aware selection was repeated 400 times using a single NumPy generator (seed 0) advanced continuously across draws, and the resulting joint-validity rates were summarized by their mean and percentile 95% interval. The retained-set curve is the primary ranking analysis; *K* = 248 is one displayed operating point.

#### Backbone geometry and novelty analyses

Backbone geometry was summarized using local distances, bond angles, absolute torsions and the O3^*′*^–P distance. For generated backbones, novelty was defined as one minus the maximum US-align TM similarity to the reference PDB library. The pair-type, binary-pair and topology-free decoders used the same 550-backbone pool; validity filtering therefore selected different subsets of this pool, whose geometry and novelty were then compared.

For the native-backbone continuous novelty analysis, each of the 94 targets was compared with its nearest match among 141 RiboCouple training-cluster representatives and the resulting US-align TM similarity was related to pair-coupled-minus-argmax changes in recovery, MCC and BP-F1. A complementary subgroup analysis reported the paired MCC and BP-F1 gains for chains with nearest-representative TM similarity at most 0.45 (*n* = 65). Spearman correlations and paired target-bootstrap confidence intervals resampled the similarity and outcome values using the same target indices. Exploratory generated-backbone stratification used the pair-type-minus-topology-free contrast.

### Statistical analysis

Confidence intervals were estimated by nonparametric bootstrap, resampling individual RNA backbones in generated-backbone analyses and RNA chains in the native-backbone benchmark. For comparisons evaluated on common backbones, paired observations were preserved by applying identical resampled unit indices to all conditions. The primary generated-backbone decoder contrasts, including comparisons among decoders on the common RNA-FrameFlow cohort, used percentile 95% confidence intervals from 2,000 bootstrap resamples (random seed 0). Paired binary joint-validity outcomes were additionally compared using exact two-sided McNemar tests, with discordant-pair counts retained in the source data.

For matched donor substitution, a nested bootstrap resampled recipient backbones and donor outcomes within each recipient. In the 94-chain native benchmark, paired effects were assessed using chain-level bootstrap 95% confidence intervals and two-sided Wilcoxon signed-rank tests. Win counts denote chains with a strictly positive pair-coupled-minus-argmax difference. To account for dependence among chains from the same structure, we additionally performed a cluster bootstrap over the 69 original PDB entries and a PDB-entry-level sign-permutation test. Four of these entries contributed 29 of the 94 chains.

The paired TM-score win rate was defined as the fraction of targets for which the reference condition achieved a higher TM-score than the comparator. Results from different structure predictors were analysed separately. Unless otherwise specified, 95% confidence intervals were two-sided.

## Data and code availability

Training and inference code, trained model parameters, evaluation scripts and the environment specification are available at https://github.com/Jakkiabc/RiboCouple. Supplementary Data 1–3 provide the 94-chain evaluation manifest, checkpoint training-data cutoffs and per-chain sequence-distant realization counts with functional-family assignments, respectively.

## Supplementary Information

## 1 Supplementary Methods

### 1.1 Data and RNA structural representations

#### 1.1.1 RNA structural data and residue indexing

Training structures were obtained from the 31 October 2023 RNA-Solo snapshot [33], comprising cleaned PDB-derived RNA chains resolved at 4.0 Å or better. We retained single RNA chains of 40–150 nucleotides and applied the structural and annotation filters described below, yielding 5,038 training examples. During data loading, structures were converted into residue-indexed rigid frames (rotations and translations), torsion representations and the node and pair features required by the flow-matching trunk. The fixed-backbone inverse-folding evaluation set was maintained separately from the training set. Dataset temporal separation was assessed using first-release dates and PDB-chain overlap with released training or evaluation manifests of RiboCouple, gRNAde, RDesign and RiboDiffusion. Training-pool overlap with the RNA-Solo source pool was also quantified for external models where released manifests permitted this comparison.

Chain length, interaction density and pseudoknot prevalence are summarized in Supplementary Table S1.

**Supplementary Table S1.** Structural composition of the 40–150-nt training window.

| Statistic | Value |
| --- | --- |
| RNA chains | 5,038 |
| Length, mean / median (nt) | 87.5 / 77 |
| Annotated interactions per chain, mean / median / maximum | 39.0 / 34 / 79 |
| Annotated-interaction density, mean / median | 0.864 / 0.894 |
| Annotated-interaction density, Q1 / Q3 / maximum | 0.795 / 1.000 / 1.350 |
| Chains with interaction density > 1 | 1,173 (23.3%) |
| Chains containing a pseudoknot | 1,801 (35.7%) |

The interaction-density statistic in Table S1 is twice the number of annotated residue pairs divided by chain length, that is, the number of paired residue endpoints per nucleotide; it can therefore exceed one.

#### 1.1.2 Residue rigid frames and torsional degrees of freedom

Let *X*_*i,a*_ *∈* ℝ^3^ denote the coordinate of atom *a* in residue *i*. Each nucleotide was represented by a residue frame

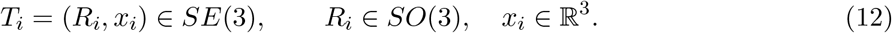

The frame origin was the C4^*′*^ coordinate, *x*_*i*_ = *X*_*i*,C4_^*′*^ . To define its orientation, we formed

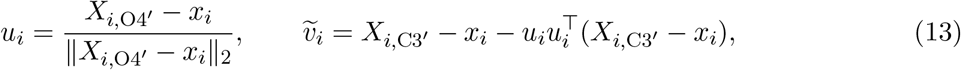

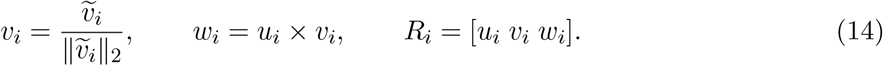

The translation locates the nucleotide, whereas the rotation orients its local sugar–backbone coordinate system. This representation separates global rigid motion from the relative placement of nucleotides.

A residue frame does not determine atoms connected through rotatable bonds. The model therefore predicts eight torsional degrees of freedom per residue as unit-circle vectors *τ*_*i,k*_ = (sin *θ*_*i,k*_, cos *θ*_*i,k*_). Frames determine residue placement and orientation; torsions determine the relative configuration of the attached rigid groups. Atomic reconstruction is described below.

RiboCouple represents an RNA of length *L* as a collection of residue frames in *SE*(3)^*L*^, with nucleotide-specific torsions completing the local backbone geometry. An invariant-point-attention (IPA) trunk updates residue representations and rigid states equivariantly, while the interaction and sequence pathways introduced below operate on the resulting node and pair representations.

#### 1.1.3 RNA rigid-group hierarchy

A rigid group is a set of atoms whose coordinates are fixed relative to a local frame. Instead of predicting every atom independently, the reconstruction represents flexibility by rotations between connected groups. The hierarchy starts from one principal residue frame and then branches into local sugar atoms, the O5^*′*^–P phosphate pathway and the C1^*′*^–N9/N1 glycosidic pathway.

The implementation contains 11 rigid-group slots: the principal residue frame and dependent groups associated with C2^*′*^, C1^*′*^, N9/N1, O3^*′*^, O5^*′*^, P, OP1, OP2, O2^*′*^ and the base/chi group. These are not independently generated global frames. The principal residue frame fixes the global placement of the nucleotide, while eight predicted torsions orient the dependent groups through this fixed kinematic hierarchy. For group *g* with parent *p*(*g*),

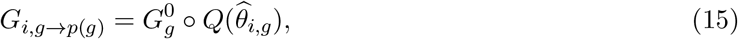

where 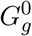 is a fixed default transform from the RNA reconstruction template and 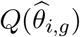 is the rotation constructed from the predicted sine–cosine pair. Groups connected through multiple bonds are transformed to the principal backbone frame by composing the transforms along their parent path,

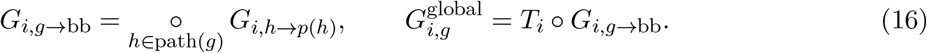

For example, the phosphate branch is composed along O5^*′*^–P– OP1, OP2, while the glycosidic branch follows C1^*′*^–N9/N1–base/chi and the O2^*′*^ branch is anchored through C2^*′*^. The neural network therefore supplies residue placement and torsional configuration, while the fixed RNA template supplies the within-group geometry. During backbone generation, reconstruction does not require a decoded nucleotide identity; the resulting coordinates are consequently a sparse backbone representation rather than a complete nucleotide-specific all-atom model.

#### 1.1.4 SSR-derived base-pair annotations

Base-pair supervision was constructed from experimental RNA structures using DSSR [34]. DSSR-annotated residue pairs were converted into binary pair matrices aligned to the model residue indices and stored with the structural examples for training. This choice follows established geometric classification of RNA base pairs [25, 26, 43, 44]; secondary-structure resources and predictors provide related but not identical pair definitions [45–48].

For DSSR-annotated base pairs, conditional pair-type labels were generated from the native nucleotide identities. Unordered AU, GC and GU combinations were assigned to three explicit classes. OTHER comprised annotated base-pair edges between valid A, C, G or U residues whose nucleotide combination did not map to AU, GC or GU. Thymine, modified residues and other unsupported residue identities were excluded from the sequence and pair-type objectives.

After symmetrization and retention of one entry per unordered residue pair (*i < j*), the 5,038-chain training subset contained 196,555 annotated base-pair edges (Supplementary Table S2). GC was the most frequent class (48.1%), followed by OTHER (23.9%), AU (19.2%) and GU (8.9%).

**Supplementary Table S2.** DSSR-derived base-pair supervision in the 5,038-chain training subset.

| Pair Type | Count | Fraction (%) |
| --- | --- | --- |
| GC | 94,499 | 48.1 |
| AU | 37,710 | 19.2 |
| GU | 17,429 | 8.9 |
| OTHER | 46,917 | 23.9 |
| Total | 196,555 | 100.0 |

### 1.2 Structural generation, relational modeling and training

#### 1.2.1 Flow matching on residue-frame space

Following flow matching [11, 35, 36], the continuous structural state of an RNA of length *L* is **T** = (*T*_1_, …, *T*_*L*_) *∈ SE*(3)^*L*^. Reference frames **T**_0_ = (**R**_0_, **x**_0_) were connected to data frames **T**_1_ = (**R**_1_, **x**_1_) by the translation and rotation interpolation paths defined in the main Methods. The training translation reference was obtained from chain-centred Gaussian coordinates and matched to data structures by batch optimal transport before interpolation. For rotations, a training reference was constructed by right-multiplying each data rotation by isotropic *SO*(3) noise sampled from the IGSO(3) distribution with *σ* = 1.5, and the intermediate rotation followed the geodesic joining the reference and data rotations. Together they define a generative transport between probability distributions rather than a physical RNA-folding trajectory. At inference, translations were initialized from the chain-centred Gaussian reference and rotations independently from the uniform distribution on *SO*(3).

Given (**T**_*t*_, *t*), the network predicts terminal frames 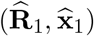. Training minimizes endpoint errors normalized by the remaining interpolation time:

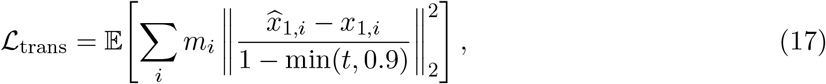

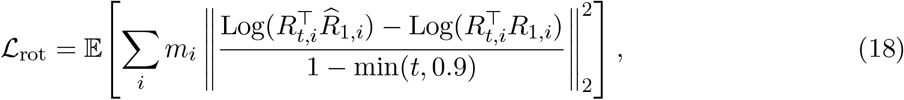

where *m*_*i*_ masks invalid or padded residues. Clipping the denominator at *t* = 0.9 prevents numerical amplification near the data endpoint. Auxiliary losses compare reconstructed frame atoms, inter-residue distances and torsion sine–cosine vectors with their data values.

The time-dependent flow state contains translations and rotations. Torsions are predicted from the trunk representation for reconstruction and auxiliary supervision and are not propagated through a separate differential equation.

At inference, the translation and rotation updates induced by the predicted endpoints were integrated for 50 steps. Sampling used a linear translation schedule and an exponential rotation schedule with rate 10: each Euler step advanced the geodesic interpolation parameter toward the predicted terminal rotation by 10 Δ*t*, so the residual rotation angle decayed exponentially across steps.

#### 1.2.2 Node, pair and geometric representations

At time *t*, the trunk maintains residue states *s*_*i*_ *∈* ℝ^256^, pair states *z*_*ij*_ *∈* ℝ^128^ and frames *T*_*i*_ *∈ SE*(3). Residue states encode flow time and residue position, whereas pair states combine sequence-separation, inter-residue geometry and self-conditioning features from a previous terminal-structure estimate. Here, *s*_*i*_ represents a residue, *z*_*ij*_ represents a directed residue–residue relation and *T*_*i*_ supplies the current three-dimensional reference system. Residue masks identify valid positions and are applied consistently to attention and geometric objectives.

#### 1.2.3 Invariant point attention and rigid-state updates

The structural trunk contains six invariant-point-attention (IPA) blocks [37]. For attention head *h*, scalar queries and keys are obtained from the residue states, pair states provide an additive relation bias, and learned query/key points are transformed from residue-local coordinates into the current global frame. The attention logit has the form

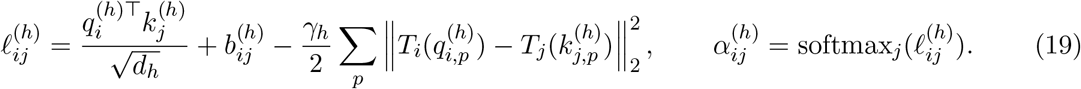

The three terms encode residue-state compatibility, the learned pair relation and the current three-dimensional placement. Scalar and point values are aggregated using *α*^(*h*)^, and point outputs are returned to the receiving residue’s local frame. This yields invariant attention weights and equivariant structural updates. Here 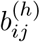 denotes the pair-derived bias used within the structural IPA trunk; the sequence decoder receives separate pair and pair-type biases defined below.

Each block applies IPA, sequence-axis Transformer processing [38] and transition layers to update *s*_*i*_. A rigid-update head maps the resulting residue state to a three-component quaternion update and a local translation,

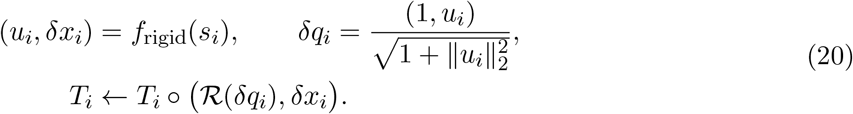

Here *ℛ* maps a unit quaternion to its rotation matrix; the local translation is rotated by the current residue frame before being added to the global position. The within-block rigid update in Eq. 20 is distinct from the rotation integration used by the flow sampler. The IPA module uses eight heads, scalar hidden dimension 128, eight query/key points and twelve value points. After six blocks, the trunk predicts the terminal frames and torsions used by the flow field and atomic reconstruction.

#### 1.2.4 Pair-representation coupling

##### Binary pair and conditional pair-type prediction

The interaction interface predicts two quantities for each residue pair: whether the residues are paired and, conditional on pairing, their nucleotide-pair class. It factorizes this relation as

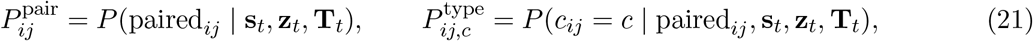

where *c ∈ {*AU, GC, GU, OTHER*}*. The binary head predicted a directed interaction logit 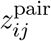 for each valid, non-diagonal residue pair. For analyses requiring an undirected representation, we symmetrized the predicted pair probabilities as

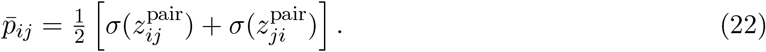

The binary objective was a masked, class-balanced binary cross-entropy over valid, non-diagonal residue pairs. Diagonal and padded positions were excluded.

At positive pair positions, the type head predicted

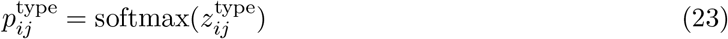

over AU, GC, GU and OTHER. Nucleotide classes were indexed in the order A, C, G and U. Supervision was restricted to annotated positive pair positions, with class balancing applied across the observed pair-type classes.

##### Pair recycling

The model applies the structural trunk twice. The first pass maps the current noised frames to residue and pair representations and predicts a soft binary interaction map,

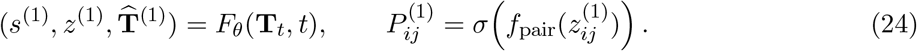

The soft probabilities are embedded into the pair pathway,

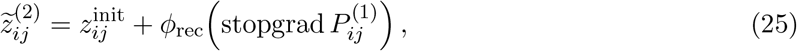

followed by

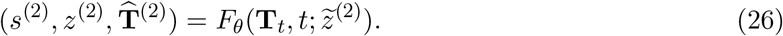

The two passes share parameters, and gradients from the second pass are not propagated through 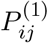. No thresholding is applied before recycling. The second-pass representations parameterize the reported pair predictions, structural endpoint and nucleotide decoder.

##### Pair-biased Transformer sequence decoder

The sequence head received residue representations from the second structural pass. As defined in the main Methods, the scalar binary pair logit and the four-dimensional conditional pair-type distribution weighted by binary pairing probability entered separate learned, head-specific attention-bias projections. These projections were applied before the padding mask. The decoder retained the continuous pair distributions and produced a categorical distribution over A, C, G and U at each valid position.

During stage-2 sequence-head training, the structural trunk and binary interaction head were frozen. The pair-type, binary-pair and topology-free sequence heads were trained independently under their respective attention-bias configurations; the pair-type head was optimized only for the pair-type condition.

#### 1.2.5 Atomic reconstruction and PDB serialization

The terminal flow state provides one principal rigid frame *T*_*i*_ = (*R*_*i*_, *x*_*i*_) per residue, and the torsion head specifies rotations between connected residue-local rigid groups. If atom *a* belongs to rigid group *g*(*a*) and has ideal within-group coordinate 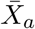, its global coordinate is

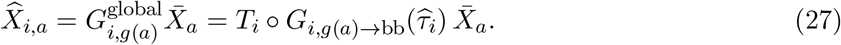

Equivalently, if the composed internal transform contains rotation *R*_*i,g*_ and translation *t*_*i,g*_,

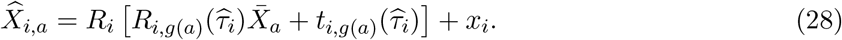

Here, *g*(*a*) denotes the rigid group containing atom *a*, and *G*_*i,g*(*a*)*→*bb_ denotes the torsion-dependent transform from that group-local frame to the residue backbone frame; *T*_*i*_ then places the residue frame in global coordinates. The model therefore predicts residue-frame placement and internal torsional configuration rather than independent Cartesian coordinates. Applying the composed rigid transforms to residue-local templates maps each template atom into global coordinates, reconstructing the sugar–phosphate backbone and the N9/N1 base anchor used for structural evaluation and PDB output. The structural output is therefore a sparse RNA backbone representation rather than a complete nucleotide all-atom model.

#### 1.2.6 Training objective and optimization

Stage 1 optimized the structural flow, auxiliary structural objectives and binary pair objective. Stage 2 initialized each sequence-decoding condition from the resulting pair-recycled structural checkpoint, froze the structural trunk and binary interaction head, and optimized the sequence head under its corresponding attention-bias configuration; the pair-type head was optimized only for the pair-type condition. The auxiliary structural atom set contained the frame atoms only; no non-frame atoms were included. A checkpoint was saved after every epoch.

#### 1.2.7 Backbone generation and nucleotide inference

Final rigid states and predicted torsions were applied to the fixed RNA reconstruction template hierarchy, and the retained backbone atoms and glycosidic anchor were serialized to PDB. After the second structural pass, residue-wise argmax over the A/C/G/U logits produced the primary sequence, which was written separately in FASTA format. One decoded sequence was evaluated per backbone in the primary benchmark.

### 1.3 Generated-backbone evaluation protocols

The shared 550-backbone cohort contained 50 backbones at each target length from 50 to 150 nucleotides in increments of 10. RhoFold+ was the primary refolding route. The primary generated-backbone benchmark and matched-adversarial analysis used TMscore-derived TM-score and RMSD [39], whereas representation-intervention and recycling analyses used US-align-derived TM-score [49] together with the corresponding TMscore-derived RMSD field. In every cohort, joint validity required both a TM-score at least 0.45 from the specified program and RMSD at most 5.5 Å from the corresponding protocol.

The RNA-FrameFlow plus gRNAde baseline used independently generated RNA-FrameFlow backbones. For the expected single-sample statistic, eight gRNAde sequences were sampled per backbone on both the RNA-FrameFlow and RiboCouple backbone pools and evaluated separately; joint validity was first summarized across backbones for each of the eight fixed sample indices, and the expected single-sample rate was computed as the mean of these eight index-specific rates, without selecting the best candidate. Multi-sample summaries under increasing sampling budgets are reported separately in Supplementary Table S5b, and frozen RiboCouple decoding on the same RNA-FrameFlow backbones is reported in Supplementary Table S5c.

The independent RFDpoly-based workflow comparison used the released RNA-only checkpoint, with 50 designs at each length from 50 to 150 nucleotides. The final sequence outputs from this workflow were refolded with RhoFold+ and evaluated with the same TMscore and joint-validity criteria used for the RiboCouple de novo comparison.

### 1.4 Evaluation cohorts and sample units

Supplementary Table S2b records the denominator and matching unit for each analysis. Cohorts from distinct frozen evaluations were analysed separately. For the pair-coupled versus argmax comparison, 10,000 paired bootstrap resamples were used; two-sided Wilcoxon signed-rank tests [50] were additionally reported for continuous paired metrics.

**Supplementary Table S2b.** Cohorts used for the principal and supplementary analyses.

| Analysis | Number | Statistical unit or matching rule |
| --- | --- | --- |
| Primary generated-backbone benchmark | 550 | Shared generated backbone |
| Predicted-to-realized pair-graph agreement | 549 | Evaluable refolded design |
| Longer-than-90-nt pair-graph analysis | 299 | Length-stratified design |
| Soft-guidance / deterministic-constraint comparison | 110 | Separately sampled matched generated backbone |
| Matched donor substitution | 110 | Separately sampled recipient backbone; four independently decoded donors per recipient |
| Interpolation and strength/type interventions | 110 | Matched generated backbone |
| Confidence-ranked edge masking | 110 | Matched generated backbone |
| Adversarial pair-graph retention | 60 | Backbone evaluated under three rewiring seeds |
| Native-backbone sequence design | 94 | Temporally held-out experimental RNA chain |
| Pair-ranking and calibration recycling analysis | 94 | Matched native RNA chain |

**Supplementary Table S2c.**
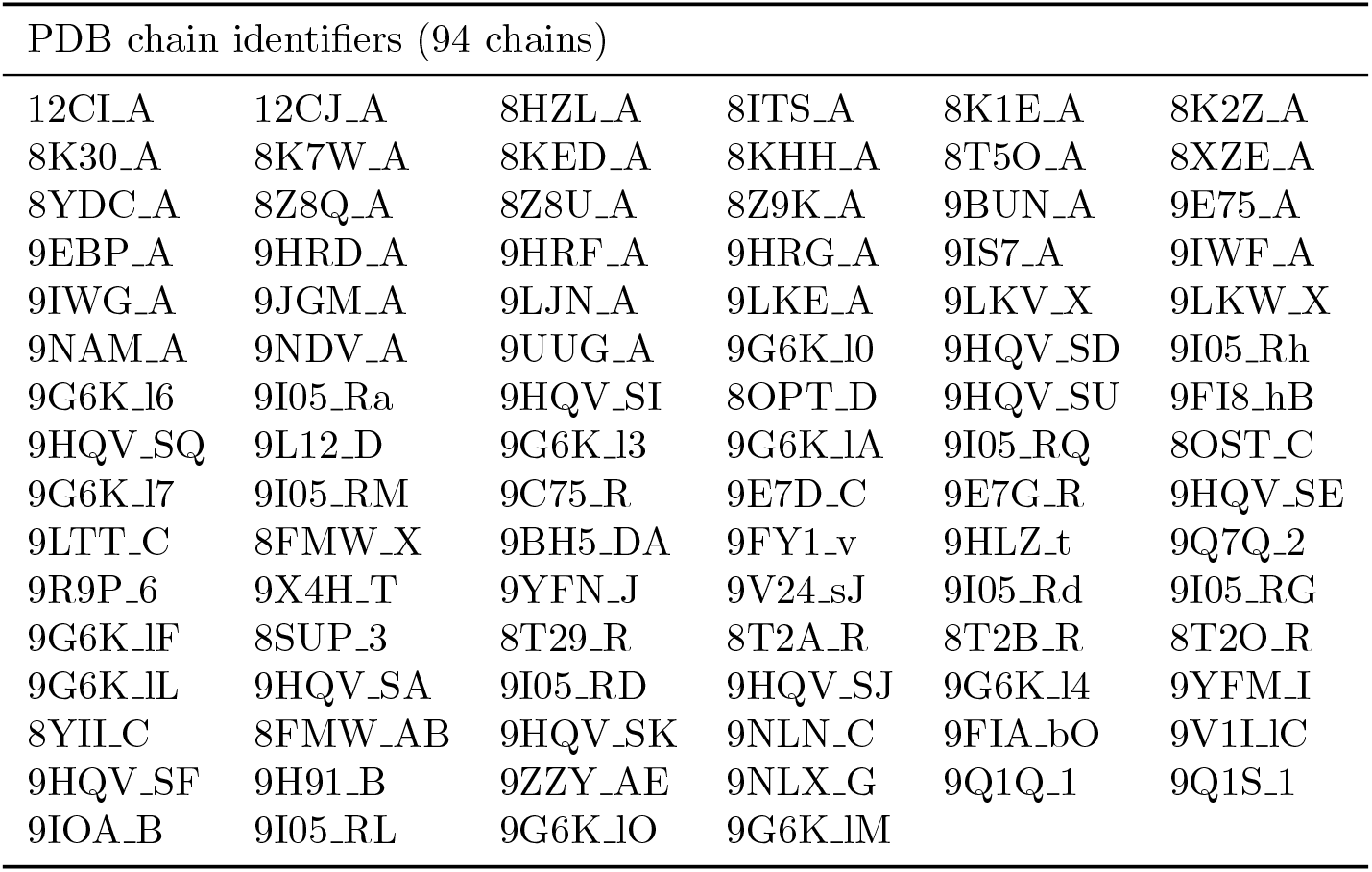
PDB chain identifiers for the 94-chain native-backbone benchmark. Chain identifiers follow the common 94-chain evaluation manifest; some multi-character labels were normalized without changing atomic coordinates.

### 1.5 Pair-graph realization and adjusted associations

#### 1.5.1 Predicted-to-realized pair-graph agreement

For the realization scatter, symmetrized pair probabilities were thresholded at *p >* 0.5 and compared with hydrogen-bonded residue pairs extracted after refolding; one design with no predicted edge above this threshold was excluded (*n* = 549). The adjusted analysis used a PairF1 covariate recomputed in a separate forward pass for all 550 backbones by comparing the predicted pair graph with the corresponding refolded pair graph. For the length-adjusted analysis, RNA length was regressed separately from realized pair-graph F1 and TM-score and Pearson correlation was calculated between the two residuals. A separate Pearson correlation was calculated within designs longer than 90 nucleotides.

#### 1.5.2 Adjusted PairF1–TM-score association

The complementary multivariable linear regression was fitted to the 550-target covariate table:

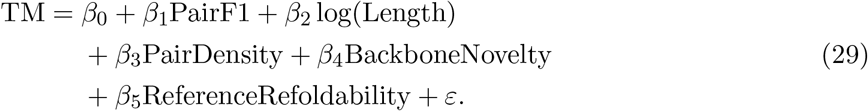

Coefficients were estimated at the target level, with nonparametric bootstrap confidence intervals. PairF1 was recomputed from a separate forward pass for all 550 targets, and its adjusted association with TM-score was estimated by regression.

### 1.6 Controlled pair-state interventions

#### 1.6.1 Matched pair-graph rewiring

Matched-adversarial graphs were constructed by degree-preserving double-edge swaps with separation-aware matching and subsequent quality control. Proposed swaps that created self-loops, duplicate edges or invalid-residue connections were rejected, and candidate swaps were constrained to retain a closely matched distribution of sequence separations |*i−j*|. Original pair strengths were reassigned to accepted adversarial edges and injected at the sequence-head interface. Three independent rewiring seeds were used. Construction quality was assessed using degree equality, retained pair count, edge overlap and separation-distribution divergence.

#### 1.6.2 Partial partner-rewiring corruption sensitivity

Sensitivity to partial loss of partner identity was evaluated in a separate 110-backbone cohort at nominal partner-rewiring corruption levels of 0%, 1%, 5%, 25%, 75% and 100% (labels c0–c100 in Extended Data Fig. 5); the achieved fraction was assessed separately. Three independent rewiring seeds were used per level. Model parameters, backbone coordinates and decoding protocol were fixed across conditions, and seed-specific outcomes were retained for the target-level summaries.

#### 1.6.3 Adversarial pair-graph retention

On a subset of 60 backbones evaluated with three rewiring seeds, the adversarial pair graph was compared with hydrogen-bonded pairs in the refolded structures. Precision, recall and F1 were calculated independently for each backbone–seed combination and then summarized across the subset.

#### 1.6.4 Type-realization accuracy for component interventions

Type-realization accuracy for the component-intervention cohort was computed as the fraction of evaluated predicted pair entries whose realized nucleotide-pair class agreed with the corresponding tested class assignment under the fixed evaluation protocol. For the strength-fixed (*p* = 0.5) condition, pair-type predictions were taken from the corresponding strength-unperturbed model forward pass and compared with canonical pair classes realized after ViennaRNA folding; target-level accuracies were averaged across the 109 targets with evaluable pair-type realization labels in this 110-backbone cohort.

#### 1.6.5 Soft attention guidance, deterministic pair constraints and pair-signal removal

On 110 matched generated backbones, four decoding strategies were evaluated with identical backbone coordinates, model parameters and decoding seeds. The full-soft condition used binary and typed pair-derived attention guidance. The confidence-gated condition retained soft pair-derived guidance only for edges with probability at least 0.95 and removed below-threshold binary and typed contributions. The true-zero condition removed pair-derived attention guidance entirely. In the deterministic condition, canonical-pair constraints were applied to the sequence after full-soft decoding.

### 1.7 Calibration, native evaluation and candidate prioritization

#### 1.7.1 Pair-probability reliability calibration

Pair-ranking and calibration diagnostics were evaluated on the 94-chain native benchmark, comprising 377,234 valid, undirected residue pairs (*i < j*). Pair-ranking and calibration used hydrogen-bond-based pair annotations, whereas native-backbone MCC and BP-F1 used canonical X3DNA-DSSR-derived reference pairs. First- and second-pass pair probabilities were compared with the corresponding binary interaction labels. Expected calibration error (ECE), all-edge Brier score and target-macro Brier score quantified probability calibration, whereas target-macro area under the precision–recall curve (AUPRC), best F1 and precision at specified recall levels quantified pair discrimination. Paired uncertainty was estimated by target-level bootstrap.

#### 1.7.2 Native-backbone sequence design

The native-backbone benchmark comprises 94 experimentally determined RNA chains with first public release dates after the compared checkpoints’ training-data cutoffs. The common chain identifiers and temporal metadata are recorded in Supplementary Table S2c and the evaluation manifest; all methods in Supplementary Table S9 use the same 94-chain cohort.

The main 94-chain benchmark used one sequence per target for every method; gRNAde results use the first sampled candidate of its 16-sample inference protocol (a fixed sample index; top-1). Secondary-structure Matthews correlation coefficient (MCC) was calculated over all unordered residue pairs (*i, j*), *i < j*, for each target [51]. Canonical reference pairs were derived from X3DNA-DSSR; the corresponding positive predicted pairs came from ViennaRNA MFE secondary-structure predictions. Pseudoknotted reference contacts that cannot be represented by the RNAfold MFE prediction were excluded from the reference pair sets for this comparison. All other residue pairs were treated as negatives. MCC therefore evaluates contact presence or absence over the full residue-pair space, whereas canonical BP-F1 evaluates overlap of the positive canonical-pair sets.

#### 1.7.3 Pair-coupled decoding rule

For each native backbone, directed binary pair probabilities were symmetrized by their arithmetic mean. Candidate edges were restricted to residues separated by at least four positions. Each residue selected its highest-probability candidate partner; an edge (*i, j*) was retained only when *i* and *j* selected one another, *i < j*, and its symmetrized probability was at least 0.8. Mutual-best selection makes retained edges non-overlapping. Decoding started from residue-wise argmax over the A/C/G/U logits. For each retained edge, the two residues were jointly reassigned to the highest-scoring member of *{*AU, UA, GC, CG, GU, UG*}*, maximizing the sum of their log-softmax nucleotide probabilities. Unselected residues retained their argmax assignments. The same fixed rule was applied to every native target.

#### 1.7.4 Pair-based candidate prioritization

Candidate prioritization used the same pair-confidence score and tie-aware cutoff rule as the main Methods. When a score tie crossed a retained-set boundary, targets strictly above the boundary were retained and the remaining positions were sampled uniformly from the tied group. Four hundred draws with seed 0 yielded the reported mean joint-validity fraction and percentile 95% interval. The plotted analysis reports retained-set sizes of 50 or greater.

### 1.8 Runtime measurement protocol

Runtime was measured after warm-up on a single NVIDIA RTX 4090 using PyTorch 2.0.1, FP32 arithmetic, and a batch size of 1 for a 100-nt RNA. Fifteen repeated measurements were summarized by the median. The structural-flow timing comprised 50 pair-recycled trunk evaluations and therefore included pair prediction and recycling at every flow step. Final decoding was timed as one additional complete forward pass comprising the trunk, pair-recycling pathway and sequence head. Checkpoint loading and external structure refolding were excluded from the reported RiboCouple timing.

## 2 Supplementary Results

*Table organization*. Supplementary Tables S3–S5d report generated-backbone comparisons and cross-generator transfer; S6–S8b document pair-graph agreement, perturbations and recycling; S9–S9e describe the 94-chain native-backbone analysis; S9f–S9g provide decoder-architecture and candidate-ranking analyses; and S10–S10b document runtime and implementation settings. Table S2b lists the denominators and matching units for each analysis.

### 2.1 Generated-backbone benchmark results

These tables cover three aspects of the generated-backbone evaluation. Supplementary Tables S3 and S3b report same-backbone decoder comparisons and sensitivity to validity thresholds; Tables S4 and S4b examine evaluator and length dependence; and Tables S5–S5d compare independent workflows, sampling protocols and transfer of frozen decoders.

**Supplementary Table S3.** Same-backbone generated-RNA sequence-design benchmark. All rows use the same 550 RiboCouple-generated backbones. The output column reports the number and selection of sequences evaluated per backbone.

| Sequence Method | Condition / Interface | Joint Validity ↑ | Output Semantics |
| --- | --- | --- | --- |
| RiboCouple pair-type | typed pair state | 45.09% | one deterministic output |
| RiboCouple binary-pair | binary pair state | 41.27% | one deterministic output |
| RiboCouple topology-free | no pair-derived attention bias | 33.45% | one deterministic output |
| gRNAd | external inverse folding | 31.80% | expected single sample |
| RIDER | external inverse folding | 26.00% | one sampled output |

**Supplementary Table S3b.**
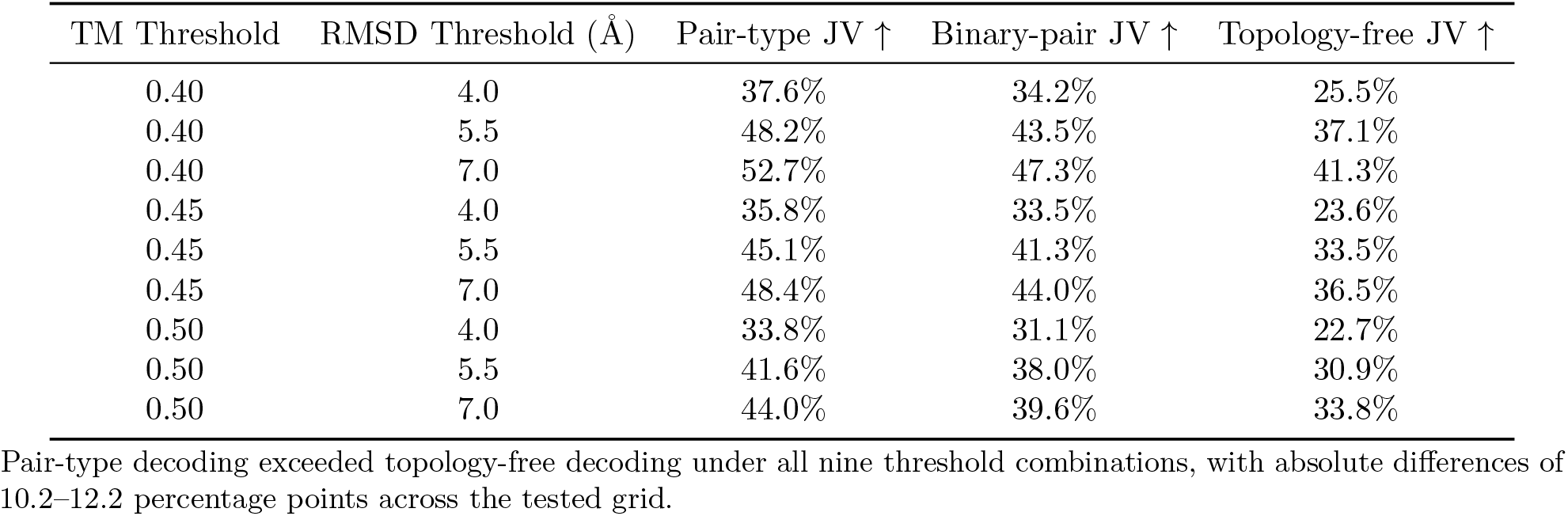
Sensitivity of joint validity to TM-score and RMSD thresholds on 550 shared backbones.

| TM Threshold | RMSD Threshold ($\text{\AA}$) | Pair-type JV ↑ | Binary-pair JV ↑ | Topology-free JV ↑ |
| --- | --- | --- | --- | --- |
| 0.40 | 4.0 | 37.6% | 34.2% | 25.5% |
| 0.40 | 5.5 | 48.2% | 43.5% | 37.1% |
| 0.40 | 7.0 | 52.7% | 47.3% | 41.3% |
| 0.45 | 4.0 | 35.8% | 33.5% | 23.6% |
| 0.45 | 5.5 | 45.1% | 41.3% | 33.5% |
| 0.45 | 7.0 | 48.4% | 44.0% | 36.5% |
| 0.50 | 4.0 | 33.8% | 31.1% | 22.7% |
| 0.50 | 5.5 | 41.6% | 38.0% | 30.9% |
| 0.50 | 7.0 | 44.0% | 39.6% | 33.8% |

Pair-type decoding exceeded topology-free decoding under all nine threshold combinations, with absolute differences of 10.2–12.2 percentage points across the tested grid.

**Supplementary Table S4.** Joint validity across structure evaluators.

| Evaluator | Pair-type ↑ | Binary-pair ↑ | Topology-free ↑ | FrameFlow + gRNAd ↑ |
| --- | --- | --- | --- | --- |
| RhoFold+ | 45.09% | 41.27% | 33.45% | 32.0% |
| Boltz-1 | 29.09% | 28.36% | 18.91% | 28.4% |
| Boltz-2 | 31.09% | 30.18% | 22.91% | 29.5% |
| trRosettaRNA2 | 30.18% | 30.00% | 21.64% | 25.3% |

**Supplementary Table S4b.** Length-dependent design performance and pair-state diagnostics.

| Metric (JV rows in %) | 50–79 nt | 80–109 nt | 110–149 nt | 150 nt |
| --- | --- | --- | --- | --- |
| $n$ | 150 | 150 | 200 | 50 |
| AUPRC mean | 0.446 | 0.240 | 0.323 | 0.089 |
| AUPRC median | 0.527 | 0.176 | 0.331 | 0.070 |
| Strong-edge confidence | 0.904 | 0.902 | 0.889 | 0.872 |
| Pair density | 0.0253 | 0.0186 | 0.0131 | 0.0146 |
| Pair entropy | 0.025 | 0.021 | 0.016 | 0.021 |
| Sequence entropy | 1.877 | 1.907 | 1.949 | 1.863 |
| Maximum run | 5 | 5 | 5 | 8 |
| Pair-type JV | 42.7% | 30.7% | 69.0% | 0.0% |
| Binary-pair JV | 43.3% | 26.7% | 61.0% | 0.0% |
| Topology-free JV | 30.0% | 16.0% | 57.5% | 0.0% |
| Reference-sequence JV | 57.7% | 65.5% | 87.0% | 21.3% |
| Pair F1 | 0.419 | 0.247 | 0.301 | 0.056 |
| Brier | 0.021 | 0.029 | 0.016 | 0.067 |
| ECE | 0.022 | 0.030 | 0.016 | 0.068 |

Values are target-level summaries from the shared 550-backbone cohort; JV, joint validity. Reference-sequence JV was computed from groups of training-set sequences within each length stratum, evaluated with the same RhoFold+ pipeline and aggregated within stratum to provide a length-stratified evaluator reference.

**Supplementary Table S5.**
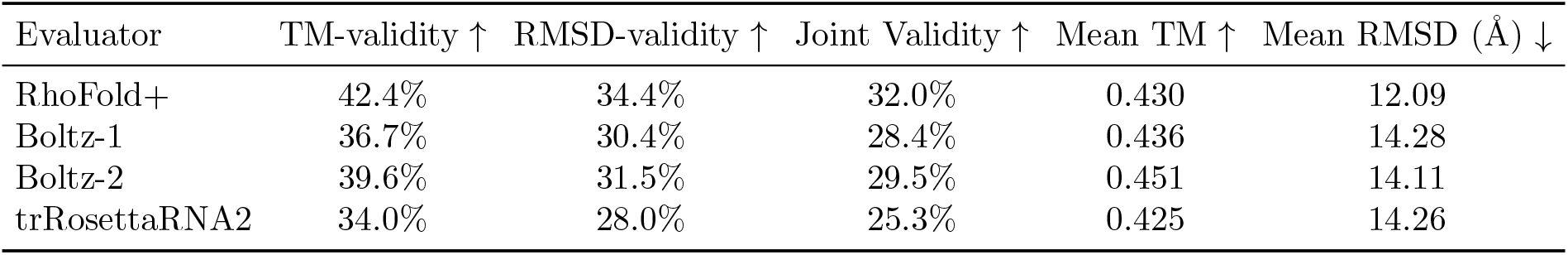
First-sampled-candidate outcomes for RNA-FrameFlow plus gRNAde on 550 generated backbones. This evaluator-specific table uses the first sampled candidate (index 0); the main-text primary statistic averages single-sample outcomes over eight fixed candidate positions.

| Evaluator | TM-validity ↑ | RMSD-validity ↑ | Joint Validity ↑ | Mean TM ↑ | Mean RMSD ($\text{\AA}$) ↓ |
| --- | --- | --- | --- | --- | --- |
| RhoFold+ | 42.4% | 34.4% | 32.0% | 0.430 | 12.09 |
| Boltz-1 | 36.7% | 30.4% | 28.4% | 0.436 | 14.28 |
| Boltz-2 | 39.6% | 31.5% | 29.5% | 0.451 | 14.11 |
| trRosettaRNA2 | 34.0% | 28.0% | 25.3% | 0.425 | 14.26 |

**Supplementary Table S5b.** gRNAde sampling-budget curve on 550 RiboCouple and 550 RNA-FrameFlow backbones.

| Pool | $K$ | Any-valid JV@$K$ ↑ | Max-TM JV@$K$ ↑ |
| --- | --- | --- | --- |
| RiboCouple | 1 | 32.00% | 32.00% |
| RiboCouple | 2 | 34.36% | 33.64% |
| RiboCouple | 4 | 38.18% | 37.82% |
| RiboCouple | 8 | 40.55% | 39.64% |
| RNA-FrameFlow | 1 | 32.00% | 32.00% |
| RNA-FrameFlow | 2 | 33.64% | 33.45% |
| RNA-FrameFlow | 4 | 35.82% | 34.91% |
| RNA-FrameFlow | 8 | 38.91% | 37.82% |

Any-valid JV@ $K$ denotes at least one joint-valid sequence among the first $K$ samples. Max-TM JV@ $K$ selects the highest-TM candidate among the first $K$ samples and records its joint validity. These multi-sample summaries are reported separately from the expected single-sample comparison used in the main text.

**Supplementary Table S5c.** Frozen RiboCouple decoding on 550 RNA-FrameFlow backbones. The same RNA-FrameFlow backbone cohort was used for all three frozen decoders.

| Decoder | Joint Validity ↑ | Mean TM ↑ |
| --- | --- | --- |
| Pair-type | 44.73% | 0.4504 |
| Binary-pair | 44.18% | 0.4514 |
| Topology-free | 37.82% | 0.4023 |

Pair-type minus topology-free joint validity was +6.91 percentage points (95% paired target-bootstrap CI, +4.18 to +10.00 percentage points; exact two-sided McNemar $P < 0.0001$ ).

**Supplementary Table S5d.** Independent length-matched de novo self-consistency comparison.

| Length (nt) | Designs | RiboCouple TM ↑ | RiboCouple JV ↑ | RFDpoly TM ↑ | RFDpoly JV ↑ |
| --- | --- | --- | --- | --- | --- |
| 50 | 50 | 0.3833 | 40.00% | 0.1469 | 0.00% |
| 60 | 50 | 0.3193 | 14.00% | 0.2174 | 0.00% |
| 70 | 50 | 0.5135 | 74.00% | 0.6459 | 98.00% |
| 80 | 50 | 0.4460 | 48.00% | 0.6857 | 100.00% |
| 90 | 50 | 0.3480 | 26.00% | 0.6123 | 92.00% |
| 100 | 50 | 0.3034 | 18.00% | 0.5286 | 30.00% |
| 110 | 50 | 0.3621 | 34.00% | 0.1395 | 0.00% |
| 120 | 50 | 0.9313 | 96.00% | 0.1103 | 0.00% |
| 130 | 50 | 0.6111 | 82.00% | 0.1063 | 0.00% |
| 140 | 50 | 0.6606 | 64.00% | 0.1005 | 0.00% |
| 150 | 50 | 0.1409 | 0.00% | 0.0950 | 0.00% |
| Overall | 550 | 0.4563 | 45.09% | 0.3080 | 29.09% |

Each workflow contributed 50 independently generated designs at each tested length. The resulting sequences were refolded with the same frozen RhoFold+ evaluator, and joint validity required TM-score at least 0.45 and RMSD at most 5.5 Å.

### 2.2 Pair-graph agreement and structural recovery

These analyses quantify agreement between predicted pair graphs and interactions in refolded structures, and the association between pair-graph agreement and structural recovery after covariate adjustment. Supplementary Table S6 reports per-design precision, recall and F1 before macro-averaging; Table S6b reports the adjusted association.

**Supplementary Table S6.** Predicted-to-realized pair-graph agreement. Precision, recall and F1 are means of per-design values.

| Quantity | Estimate | Statistical information |
| --- | --- | --- |
| Mean pair-graph precision | 0.225 | $n = 549$ designs |
| Mean pair-graph recall | 0.480 | $n = 549$ designs |
| Mean pair-graph F1 | 0.283 | $n = 549$ designs |
| F1-TM correlation | 0.607 | $P = 1.52 \times 10^{-56}$ |
| Length-adjusted correlation | 0.699 | $P = 1.37 \times 10^{-81}$ |
| Correlation, length > 90 nt | 0.830 | $n = 299, P = 2.28 \times 10^{-77}$ |

**Supplementary Table S6b.**
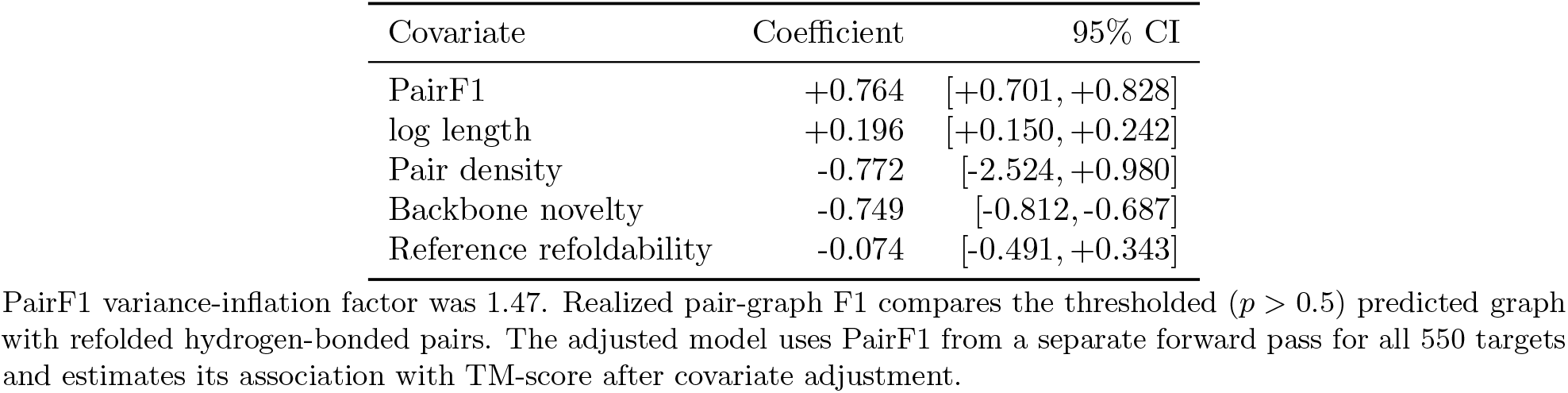
Multivariable association between predicted pair-graph agreement and TM-score. The 549-design realization scatter compares thresholded predicted pair graphs with refolded pair graphs. The adjusted coefficient uses predicted PairF1 recomputed in a separate forward pass for all 550 targets and was estimated with the target-level model in Eq. (29).

| Covariate | Coefficient | 95% CI |
| --- | --- | --- |
| PairF1 | +0.764 | [+0.701, +0.828] |
| log length | +0.196 | [+0.150, +0.242] |
| Pair density | -0.772 | [-2.524, +0.980] |
| Backbone novelty | -0.749 | [-0.812, -0.687] |
| Reference refoldability | -0.074 | [-0.491, +0.343] |

PairF1 variance-inflation factor was 1.47. Realized pair-graph F1 compares the thresholded ( $p > 0.5$ ) predicted graph with refolded hydrogen-bonded pairs. The adjusted model uses PairF1 from a separate forward pass for all 550 targets and estimates its association with TM-score after covariate adjustment.

The association remained positive after adjustment for RNA length and the measured backbone covariates.

### 2.3 Effects of pair-graph rewiring

#### Adversarial rewiring and sequence-quality controls

These analyses tested how decoding responds to the placement of pair relations under degree-matched rewiring. Double-edge swaps preserve edge count and residue degrees exactly. After quality-control filtering, final injected graphs retained 92.3% of the original pair count on average (eight-backbone construction assessment); smoothed-count KL divergence between sequence-separation distributions was at most 0.39 and residual overlap with the original graph was 15–32%. Injected adversarial edges had mean F1 of 0.018 against refolded hydrogen-bond pairs in the 60-backbone retention subset (Extended Data Fig. 3).

**Supplementary Table S7.** Matched-adversarial outcomes on 550 generated backbones.

| Condition | Joint Validity ↑ | Mean TM-score ↑ | ΔTM | Win rate ↑ |
| --- | --- | --- | --- | --- |
| Normal | 45.09% | 0.456 | – | – |
| Matched-adversarial, three seeds | 0.00% | 0.125–0.127 | 0.329–0.332 | 90.0–90.4% |

All four arms were reevaluated with RhoFold+ and TMscore on the frozen 550-target set. The Normal arm used the same argmax sequences as the primary benchmark and reproduced 248/550 valid designs. Rewired values are summarized across three intervention seeds. $\Delta$ TM is normal minus rewired; win rate is the fraction of targets with higher TM-score under Normal.

**Supplementary Table S7b.** Sequence-level quality diagnostics on 550 generated-backbone targets.

| Condition | Mean GC (%) | Median Entropy | Median Maximum Run | Run $\geq 6$ | Run $\geq 8$ |
| --- | --- | --- | --- | --- | --- |
| Normal pair-type representation | 54.9% | 1.944 | 5 | 34.9% | 15.6% |
| Matched-rewired pair representation | 47.2% | 1.788 | 7 | 68.2% | 35.6% |

Entropy is nucleotide-composition Shannon entropy; maximum run is the longest homopolymer. Normal and rewired sequence diagnostics are paired across the same 550 targets. The mean GC change (rewired minus Normal) was $-7.7$ percentage points (95% bootstrap confidence interval, $-9.0$ to $-6.5$ percentage points).

**Supplementary Table S7c.**
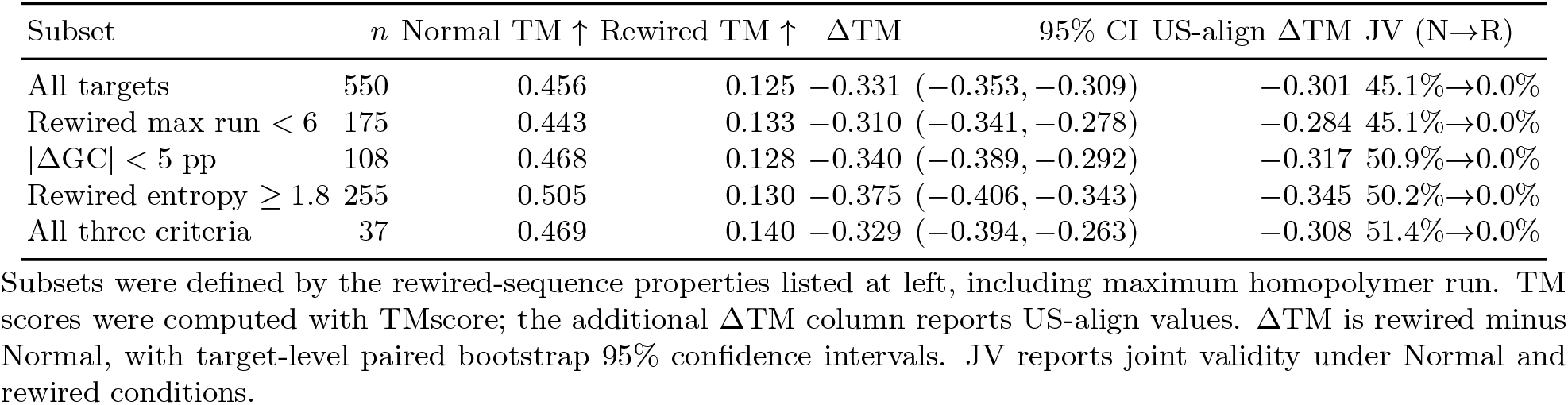
Structural robustness within rewired sequences lacking marked sequence abnormalities.

Sequence-level quality was assessed on the same 550 targets. Rewiring shifted sequence composition and increased homopolymer length, whereas structural recovery remained substantially lower after rewiring, including among designs with limited changes in GC content, nucleotide entropy or homopolymer length (Supplementary Tables S7b,c).

A complementary broad permutation reassigned complete pair-strength and conditional-class records across off-diagonal residue-pair entries in the 110-backbone intervention cohort, analyzed separately from the 550-backbone rewiring analysis in Supplementary Table S7. Joint validity decreased from 48.2% with the aligned full-soft representation to 0% after permutation, and mean TM-score decreased from 0.469 to 0.112; matched partner rewiring (joint validity 0%, mean TM-score 0.143) and a zeroed representation (joint validity 0%, mean TM-score 0.112) provided reference interventions in the same cohort. This positional-misalignment test shows strong sensitivity to the alignment of relational records with recipient residue-pair positions.

#### Matched donor substitution and interpolation

Matched donor substitution and interpolation evaluated the sensitivity of decoding to backbone alignment while controlling for global pair-state statistics (Supplementary Tables S7d,e).

**Supplementary Table S7d.** Matched donor-state substitution on 110 generated backbones.

| Condition | Joint Validity (%) ↑ | Mean TM-score ↑ |
| --- | --- | --- |
| Aligned recipient state | 46.36% | 0.4474 |
| Matched donor substitution | 21.36% | 0.3194 |
| Aligned minus donor | 25.00 pp | +0.1280 |

Each recipient was evaluated with four independently decoded, exact-length donor states and donor outcomes were averaged only after evaluation. The nested target-and-donor bootstrap 95% confidence interval for the aligned-minus-donor joint-validity difference was 16.4-33.6 percentage points. Donor matching distance had median 0.211 and 90th percentile 0.553.

Sequence-composition diagnostics were comparable between the two conditions (mean GC 54.4% versus 52.9%, nucleotide-composition Shannon entropy 1.905 versus 1.908 and mean longest homopolymer run 5.90 versus 5.81 for aligned recipient and matched donor states, respectively), indicating that donor substitution reduced structural recovery without comparably large changes in these sequence-composition measures.

**Supplementary Table S7e.** Aligned-to-donor pair-state interpolation.

| Donor Mixing Fraction $\alpha$ | Joint Validity ↑ | Mean TM-score ↑ |
| --- | --- | --- |
| 0.00 | 46.36% | 0.4474 |
| 0.25 | 46.36% | 0.4429 |
| 0.50 | 26.36% | 0.3609 |
| 0.75 | 20.91% | 0.3266 |
| 1.00 | 21.82% | 0.3209 |

Mean TM-score deteriorated progressively as the donor contribution increased. Joint validity showed the same overall transition but was not strictly monotonic.

#### Component localization and confidence-ranked masking

Pair-strength and pair-type perturbations, together with confidence-ranked masking, were used to assess decoder sensitivity to individual components of the relational state (Supplementary Tables S7f,g).

**Supplementary Table S7f.** Pair-strength and pair-type decomposition in the frozen decoder.

| Condition | Joint Validity ↑ | Mean TM-score ↑ | Type-realization accuracy ↑ |
| --- | --- | --- | --- |
| Normal | 46.36% | 0.4474 | 51.0% |
| Strength fixed ($p = 0.5$) | 22.73% | 0.3388 | 14.2% |
| Strength shuffled | 32.73% | 0.4035 | 51.8% |
| Type shuffled | 47.27% | 0.4483 | 51.4% |
| Type uniform | 47.27% | 0.4453 | 50.9% |

Acute perturbation of pair strength reduced structural recovery, whereas perturbing the explicit conditional pair-type channel produced little average change in the tested endpoints. For the strength-fixed condition, type-realization accuracy was 14.2% across 109 evaluable targets in the 110-backbone cohort, using the same target-level averaging as the other conditions. These fixed-decoder interventions are distinct from the comparison between independently trained binary-pair and pair-type models.

**Supplementary Table S7g.** Confidence-ranked masking of strong pair relations.

| Masking Condition | Joint Validity ↑ |
| --- | --- |
| None | 46.36% |
| Lowest-confidence 10% | 46.36% |
| Lowest-confidence 20% | 45.45% |
| Lowest-confidence 30% | 42.73% |
| Highest-confidence 10% | 32.73% |
| Highest-confidence 20% | 17.27% |
| Highest-confidence 30% | 9.09% |

Matched masking fractions removed strong relations from opposite ends of the confidence ranking.

A separate 110-target partial partner-rewiring corruption-sensitivity analysis evaluated nominal corruption levels of 0%, 1%, 5%, 25%, 75% and 100% (c0–c100 in Extended Data Fig. 5; three seeds per level).

### 2.4 Soft attention guidance versus deterministic pair constraints

Supplementary Table S8a asks how full-soft pair guidance compares with confidence-gated soft guidance, post-decoding deterministic pairing and removal of pair-derived guidance (Fig. 4d,e). All four conditions used the same separately sampled 110-backbone cohort, model parameters and decoding seeds.

Confidence-gated soft guidance retained intermediate performance, whereas post-decoding deterministic enforcement did not reproduce the performance of full-soft guidance in this setting.

**Supplementary Table S8a.** Four-condition pair guidance and deterministic constraint on 110 matched generated backbones.

| Condition | Joint Validity ↑ |
| --- | --- |
| Full-soft attention guidance | 48.2% |
| Soft pair-derived guidance retained at $p \geq 0.95$ | 24.5% |
| Deterministic canonical-pair enforcement after decoding | 9.1% |
| True zero pair-derived guidance | 0.0% |

The deterministic condition applies canonical-pair enforcement after soft decoding. The true-zero condition removes both binary and typed pair-derived attention guidance.

**Supplementary Table S8b.** Pair recycling.

| Metric | Pass 1 | Pass 2 | Paired change | 95% CI |
| --- | --- | --- | --- | --- |
| AUPRC ↑ | 0.649 | 0.673 | +0.024 | [+0.012, +0.035] |
| Best F1 ↑ | 0.646 | 0.651 | +0.005 | [−0.004, +0.015] |
| Precision at recall 0.5 ↑ | 0.705 | 0.740 | +0.035 | [+0.013, +0.056] |
| Precision at recall 0.8 ↑ | 0.425 | 0.391 | −0.035 | [−0.050, −0.020] |
| Pooled ECE ↓ | 0.024 | 0.013 | | Not reported |
| All-edge Brier score ↓ | 0.017 | 0.011 | | Not reported |
| Target-macro Brier score ↓ | 0.018 | 0.013 | −0.0051 | [−0.0057, −0.0045] |

*Note.* Ranking and calibration metrics were evaluated on the 94-chain native benchmark (377,234 valid undirected residue pairs), using hydrogen-bond-based pair labels, distinct from the X3DNA-DSSR-derived canonical reference used for S9 inverse-folding scores.

### 2.5 Recycling improves pair calibration and changes downstream design

These analyses distinguish whether recycling improves pair-probability ranking and calibration from whether its refined pair logits benefit downstream sequence decoding. On the 94-chain native benchmark, recycling increased AUPRC from 0.649 to 0.673 and reduced calibration errors, although precision at 80% recall declined (Supplementary Table S8b; Extended Data Fig. 4). In a fixed-decoder comparison, using first-pass instead of second-pass binary pair logits reduced joint validity by 6.4 percentage points and mean TM-score by 0.024 (Extended Data Fig. 2).

### 2.6 Native-backbone inverse-folding comparison

The native-backbone tables distinguish single-output inverse-folding benchmarks (Supplementary Table S9), paired effects and alternative pairing rules (Tables S9b and S9c), and sequence-distant pairing compatibility (Tables S9d and S9e). All methods in the 94-chain evaluation contribute one sequence per target, with full-length sequence recovery, 2D MCC and canonical BP-F1 computed under common reference and prediction conventions. Pair-coupled and residue-wise argmax decoding used identical backbone inputs and differed only in the decoding rule, whereas each external inverse-folding baseline used its released inference implementation. RIDER was evaluated with a single output per target. Supplementary Table S9c reports a separate diagnostic comparing alternative pair-constraint decoding strategies on the same 94 chains. For the sequence-distant realization analysis (Supplementary Tables S9d and S9e), native canonical pairs from the same X3DNA-DSSR reference were restricted to pairs separated by at least 10 residues in sequence, a threshold fixed before the analysis was run; realization was scored per chain as canonical compatibility, defined here to include Watson–Crick and GU wobble combinations (AU, UA, GC, CG, GU or UG) at the two natively paired positions, irrespective of the native combination, and as identity (the exact native pair), with paired target-level bootstrap intervals (10,000 resamples, seed 20261021), a PDB-entry-level clustered bootstrap and two-sided Wilcoxon signed-rank tests. Functional families were assigned from RCSB entry and entity annotations by a fixed keyword rule, independently of model outputs and realization rates.

**Supplementary Table S9.**
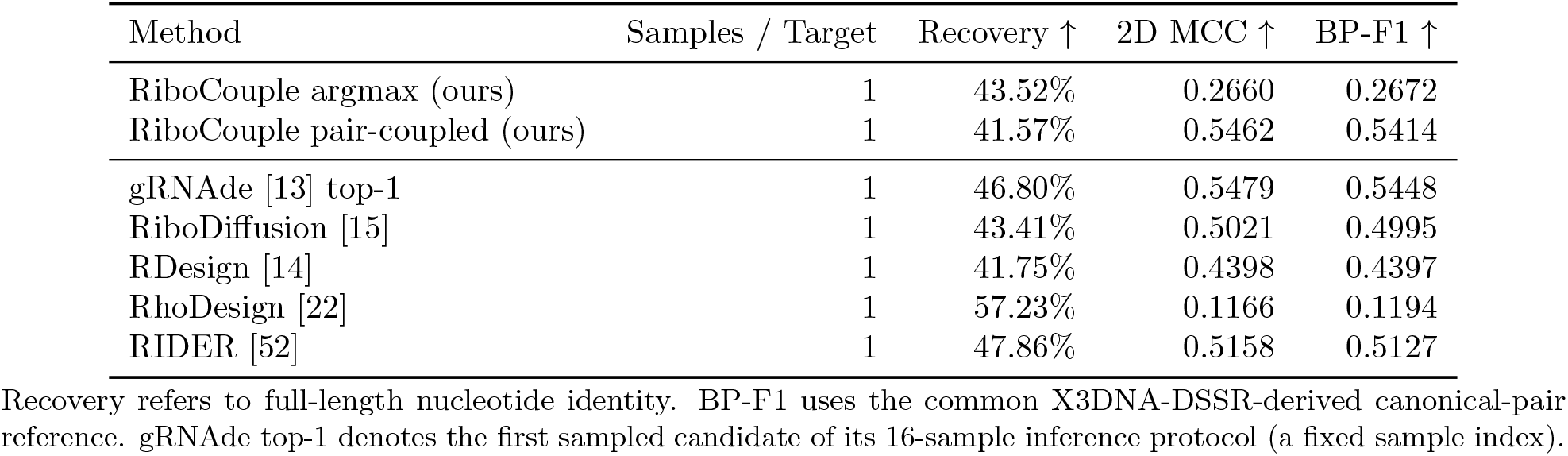
Fixed-backbone inverse folding and secondary-structure agreement on 94 temporally held-out native RNA chains. All methods use a single generated sequence per target and the same 94 target chains. Canonical BP-F1 and 2D MCC use X3DNA-DSSR-derived reference pairings and the common ViennaRNA MFE prediction protocol.

**Supplementary Table S9b.** Paired effects of pair-coupled versus argmax decoding on the same 94 native RNA chains. Intervals are target-level paired bootstrap 95% CIs; two-sided *P* values are from Wilcoxon signed-rank tests.

| Metric | Argmax | Pair-coupled | $\Delta$ | 95% CI of $\Delta$ | Wins / 94 | $P$ |
| --- | --- | --- | --- | --- | --- | --- |
| Recovery ↑ | 43.52% | 41.57% | -1.94 pp | [-2.77, -1.08] pp | 24 | $4.1 \times 10^{-5}$ |
| 2D MCC ↑ | 0.2660 | 0.5462 | +0.2802 | [0.225, 0.338] | 75 | $1.89 \times 10^{-12}$ |
| Canonical BP-F1 ↑ | 0.2672 | 0.5414 | +0.2742 | [0.220, 0.328] | 73 | $1.93 \times 10^{-12}$ |

Recovery change is expressed in percentage points (pp) and calculated from unrounded per-target values; the displayed group means were rounded independently, so subtracting the displayed percentages may differ by 0.01 pp. MCC and BP-F1 differences use absolute metric units. Wins count targets where pair-coupled exceeds argmax; ties are not counted as wins. For the MCC gain, a separate PDB-entry bootstrap across 69 source entries gave a 95% CI of [0.221, 0.345], versus the chain-level CI of [0.225, 0.338]; PDB-entry-level sign permutation gave $P \leq 10^{-5}$ .

**Supplementary Table S9c.** Alternative pair-constraint decoding strategies on the same 94 native RNA chains. Predicted-binary and native-reference-binary conditions use model-predicted and reference pair graphs, respectively. The native-reference-binary graph uses X3DNA-DSSR-derived canonical pair annotations.

| Condition | Recovery ↑ | 2D MCC ↑ | Canonical BP-F1 ↑ |
| --- | --- | --- | --- |
| Argmax | 43.52% | 0.2660 | 0.2672 |
| Pair-coupled | 41.57% | 0.5462 | 0.5414 |
| Predicted binary pairs | 41.58% | 0.5236 | 0.5195 |
| Native-reference binary pairs | 44.00% | 0.4921 | 0.4911 |

At sequence-distant native canonical pairs (separated by at least 10 residues; 1,646 pairs across all 94 chains), pair-coupled decoding increased mean per-chain canonical compatibility from 74.67% to 96.30%. Exact native-pair identity increased more modestly, from 29.92% to 37.54%, consistent with the canonical assignment vocabulary of the decoder. The canonical-compatibility gain persisted under PDB-entry-level resampling; paired differences, intervals, tests and win counts are reported in Supplementary Table S9d.

After stratification into twelve functional families using a fixed keyword rule based on RCSB entry and entity annotations, independently of model outputs and realization rates, positive pooled gains were observed in every family, including ribozymes, IRES elements and other long non-coding RNAs, whose biological functions are associated with long-range secondary-structure organization. Canonical compatibility for tRNA pairs reached 99.4%. Riboswitches and aptamers showed smaller absolute gains, consistent with residue-wise argmax already achieving 81–87% on these small, strongly constrained folds (Supplementary Table S9e). The pair-coupled joint-assignment step selects among six canonical combinations (AU, UA, GC, CG, GU and UG); noncanonical interactions can be represented by the pair-type head but are not produced by this assignment rule.

**Supplementary Table S9d.** Supplementary Table S9d. Canonical compatibility at sequence-distant native pairs on the same 94 native RNA chains.

| Metric (rates in %) | Argmax | Pair-coupled | $\Delta$ (pp) | 95% CI (pp) | $P$ | Wins |
| --- | --- | --- | --- | --- | --- | --- |
| Canonical compatibility, per-chain mean | 74.67% | 96.30% | +21.62 | [+18.1, +25.5] | $1.7 \times 10^{-15}$ | 83/94 |
| Canonical compatibility, pooled over pairs | 76.79% | 96.35% | +19.56 | — | — | — |
| Identity, per-chain mean | 29.92% | 37.54% | +7.62 | [+5.5, +9.9] | $1.9 \times 10^{-10}$ | 60/94 |
| Identity, pooled over pairs | 31.96% | 38.15% | +6.19 | — | — | — |

*Note.* Native canonical pairs separated by at least 10 residues in sequence (X3DNA-DSSR-derived reference; 1,646 pairs). Canonical compatibility requires any canonical combination (AU, UA, GC, CG, GU or UG) at the two natively paired positions, irrespective of the native combination; identity realization requires the exact native pair. Intervals are target-level paired bootstrap 95% CIs (10,000 resamples, seed 20261021); a bootstrap resampling the 69 PDB entries rather than individual chains gave a 95% CI of +16.7 to +25.4 percentage points for the canonical-compatibility difference. Two-sided $P$ values are from Wilcoxon signed-rank tests; wins count chains on which pair-coupled exceeded argmax. Inference statistics (CI, $P$ and wins) are reported for per-chain mean differences only; dashes in pooled-over-pair rows indicate that the corresponding statistics are not reported for that aggregation. Per-chain counts are provided in Supplementary Data 3.

**Supplementary Table S9e.** Family-stratified canonical compatibility at sequence-distant native pairs.

| Family | Chains | Pairs | Argmax (%) | Pair-coupled (%) | $\Delta$ (pp) | Wins |
| --- | --- | --- | --- | --- | --- | --- |
| rRNA | 32 | 304 | 76.0% | 94.7% | +18.8 | 23/32 |
| Riboswitch | 12 | 264 | 86.7% | 97.3% | +10.6 | 11/12 |
| Aptamer | 10 | 188 | 80.9% | 95.2% | +14.4 | 10/10 |
| tRNA | 9 | 173 | 80.3% | 99.4% | +19.1 | 9/9 |
| Other ncRNA | 8 | 207 | 67.1% | 93.7% | +26.6 | 7/8 |
| Viral ncRNA | 8 | 199 | 79.9% | 98.0% | +18.1 | 8/8 |
| IRES | 4 | 119 | 63.0% | 93.3% | +30.3 | 4/4 |
| Ribozyme | 4 | 48 | 72.9% | 100.0% | +27.1 | 4/4 |
| Small RNA | 3 | 59 | 66.1% | 98.3% | +32.2 | 3/3 |
| snoRNA | 2 | 47 | 80.9% | 100.0% | +19.1 | 2/2 |
| Synthetic | 1 | 29 | 75.9% | 96.6% | +20.7 | 1/1 |
| crRNA | 1 | 9 | 66.7% | 100.0% | +33.3 | 1/1 |

*Note.* Functional families were assigned from RCSB entry and entity annotations by a fixed keyword rule, independently of model outputs and realization rates. Rates are pooled over pairs within each family; $\Delta$ is the pooled pair-coupled-minus-argmax difference; wins count chains on which pair-coupled exceeded argmax. Family assignments and per-chain counts are provided in Supplementary Data 3.

### 2.7 Sequence-head architecture comparison

Four sequence-head conditions were evaluated on the same 550 backbone targets using RhoFold+, TMscore and the primary validity thresholds: the pair-type Transformer, binary-pair Transformer, topology-free Transformer and a multilayer perceptron (MLP) sequence head. The topology-free Transformer used for the primary comparison contains no pair-bias projection. Because these sequence heads were trained independently, differences among rows also reflect training history (Supplementary Table S9f).

**Supplementary Table S9f.** wAggregate evaluation of four sequence-decoding conditions on 550 shared backbone targets.

| Sequence Head | Mean TM ↑ | Mean RMSD ↓ | TM-valid ↑ | RMSD-valid ↑ | Joint-valid ↑ |
| --- | --- | --- | --- | --- | --- |
| Pair-type Transformer | 0.4563 | 11.33 | 50.18% | 50.91% | 45.09% |
| Binary-pair Transformer | 0.4485 | 11.58 | 47.45% | 47.09% | 41.27% |
| Topology-free Transformer | 0.3950 | 13.83 | 39.45% | 38.73% | 33.45% |
| MLP sequence head | 0.2596 | 19.81 | 16.18% | 18.36% | 14.91% |

All conditions used the same 550 backbone targets, RhoFold+ refolding, TM-score calculated with TMScore, TM-score threshold $\geq 0.45$ and RMSD threshold $\leq 5.5$ Å.

### 2.8 Length-dependent performance

Length-stratified joint validity was highest at 110–149 nt, lower at 50–79 and 80–109 nt, and zero for all three decoders at 150 nt (Supplementary Table S4b). The separate length-stratified reference-sequence control was also lowest at 150 nt (21.3%). The control comprised groups of training-set sequences evaluated with the same RhoFold+ pipeline. The 80–109-nt trough coincided with lower pair-head discrimination (mean AUPRC, 0.240; pair F1, 0.247), despite stable strong-edge confidence, sequence entropy and maximum run length. Backbone composition and refolding difficulty may both contribute to this length-dependent pattern.

### 2.9 Confidence-based candidate prioritization

Pair confidence was associated with downstream joint validity and was used to rank candidates before external refolding (Supplementary Table S9g). Tie-aware ranking used the sum of the ten largest binary pair probabilities over directed off-diagonal pairs (*i < j*). At *K* = 248, joint-validity rates for pair-type and topology-free decoding were 59.6% and 46.7%, respectively, compared with 45.1% and 33.5% without filtering; across *K* = 50–248, pair-type remained 12.8–13.6 percentage points above topology-free decoding. Confidence enriched pair-type outcomes across four independent structure evaluators and, after frozen RiboCouple decoding, on RFDpoly and RNA-FrameFlow backbones (Fig. 6). Target-level bootstrap intervals for the top-248 enrichment over the unfiltered rate excluded zero on all three backbone cohorts (95% CI, +10.5 to +18.6 percentage points on RiboCouple backbones, +11.8 to +19.1 on RFDpoly and +13.3 to +21.1 on RNA-FrameFlow; 2,000 length-stratified resamples, re-ranking within each resample). Across eleven 50-target length strata, within-stratum correlations between pair confidence and RhoFold+ joint validity were positive in eight of the ten informative strata (median Spearman *ρ* = 0.27; maximum *ρ* = 0.54 at 100 nt), near zero at 70 and 110 nt, and undefined at 150 nt because no designs were joint-valid. Pair confidence therefore also ranked outcomes within most lengths, with strength varying across lengths.

**Supplementary Table S9g.** Pair-head confidence stratifies downstream designability across structure evaluators.

| Evaluator | Spearman $\rho$ | AUROC ↑ | All JV ↑ | Top-248 JV ↑ | 95% tie interval | Enrichment ↑ |
| --- | --- | --- | --- | --- | --- | --- |
| RhoFold+ | 0.430 | 0.701 | 45.1% | 59.6% | [58.1%, 60.9%] | 1.32× |
| Boltz-1 | 0.358 | 0.650 | 29.1% | 37.6% | [35.9%, 39.1%] | 1.29× |
| Boltz-2 | 0.327 | 0.632 | 31.1% | 38.8% | [37.1%, 39.9%] | 1.25× |
| trRosettaRNA2 | 0.452 | 0.734 | 30.2% | 46.8% | [45.6%, 48.0%] | 1.55× |

The score is the sum of the ten largest binary pair probabilities over valid off-diagonal residue pairs, taking the directed entry with $i < j$ for each unordered pair. Spearman correlation uses average ranks for tied scores and relates this score to evaluator-specific TM-score; AUROC treats evaluator-specific joint validity as the binary outcome. For top-248 selection, targets strictly above the cutoff were retained and tied boundary targets were sampled uniformly over 400 draws (seed 0); values are means with percentile 95% intervals across tie-resolution draws. All analyses use the same 550 pair-type designs. The ranking score uses pair probabilities alone, before external refolding.

### 2.10 Generated-backbone novelty and validity-based selection

The mean nearest-PDB TM similarity was 0.589, corresponding to a mean novelty of 0.411. Novelty shifts after joint-validity filtering reflect selection of different valid subsets (Extended Data Fig. 6). In a descriptive novelty-stratified analysis, the pair-type-minus-topology-free joint-validity difference was 17.5, 14.8 and 2.7 percentage points in the low-, intermediate- and high-novelty terciles, respectively.

### 2.11 Runtime of structural generation and sequence decoding

On a 100-nt RNA evaluated with batch size 1 on a single NVIDIA RTX 4090, RiboCouple backbone generation and sequence decoding together required approximately 9.5 s; the 50-step pair-recycled structural flow dominated runtime (median, 9.31 s), whereas sequence decoding itself required 0.16 s (Supplementary Table S10). Pair-type, binary-pair and topology-free Transformer decoding operated at comparable subsecond latency. Stage-wise gRNAde measurements are included as reference timings for sequence generation from a supplied backbone. The one- and 16-candidate measurements were 6.30 and 6.25 s, respectively, because the 16 candidates were generated as a batch; multi-sample selection additionally requires evaluation of the candidate structures.

**Supplementary Table S10.**
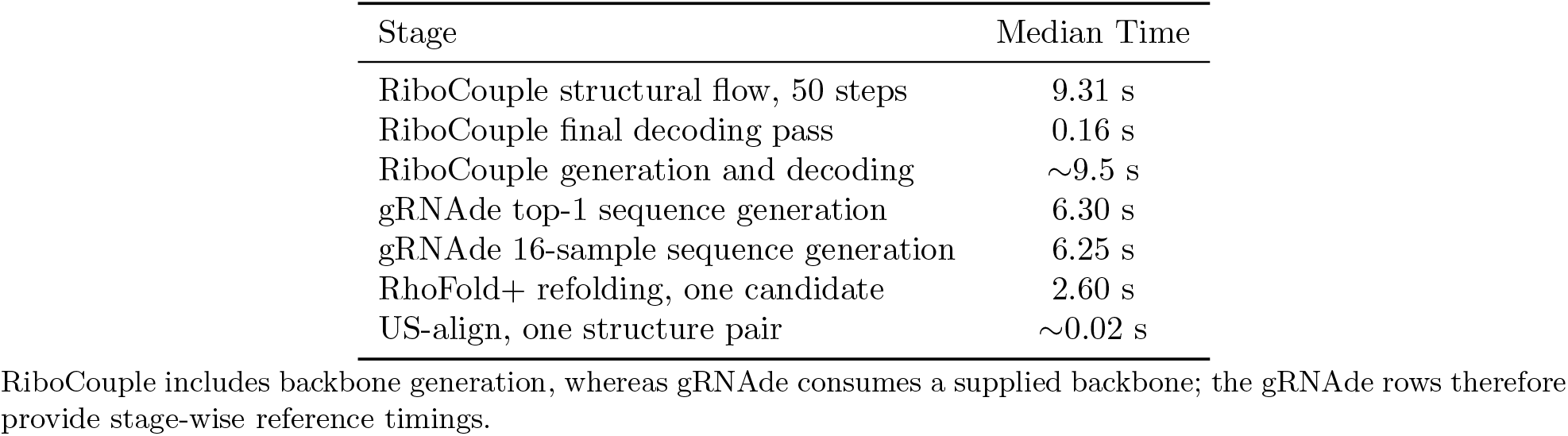
Stage-wise inference runtime on a 100-nt RNA. RiboCouple timings were measured over 15 post-warm-up runs using an NVIDIA RTX 4090, PyTorch 2.0.1, FP32 arithmetic and batch size 1.

**Supplementary Table S10b.** Training and inference configuration for RiboCouple.

| Setting | Value |
| --- | --- |
| Stage-1 binary pair-loss weight, $\lambda_{\text{pair}}$ | 1.0 |
| Stage-1 translation / rotation / auxiliary structural-loss weights | 2.0 / 1.0 / 1.0 |
| Stage-2 pair-type loss weight, $\lambda_{\text{type}}$ | 0.2 |
| Stage-2 sequence loss weight, $\lambda_{\text{seq}}$ | 1.0 |
| Stage-2 frozen parameters | Structural trunk and binary interaction head |
| Optimizer | AdamW; learning rate $10^{-4}$; weight decay 0.01; no scheduler |
| Training | Single node, 4× NVIDIA GeForce RTX 4090 (24 GB each), DDP; batch size 28 per device; at most 200 epochs |
| Checkpoint selection | TM-validity on a separate unconditional-evaluation cohort |
| Pair-coupled native decoding | Symmetrized pair probability; minimum sequence separation 4; reciprocal-best non-overlapping edges; probability threshold 0.8 |

Stage 2 initialized each independently trained sequence-decoding condition from the pair-recycled structural checkpoint; the pair-type loss was omitted for the binary-pair and topology-free conditions. The pair-coupled native-decoding rule and target manifest are provided in the accompanying release package.

## 3 Extended Data Figures

**Extended Data Figure 1.**
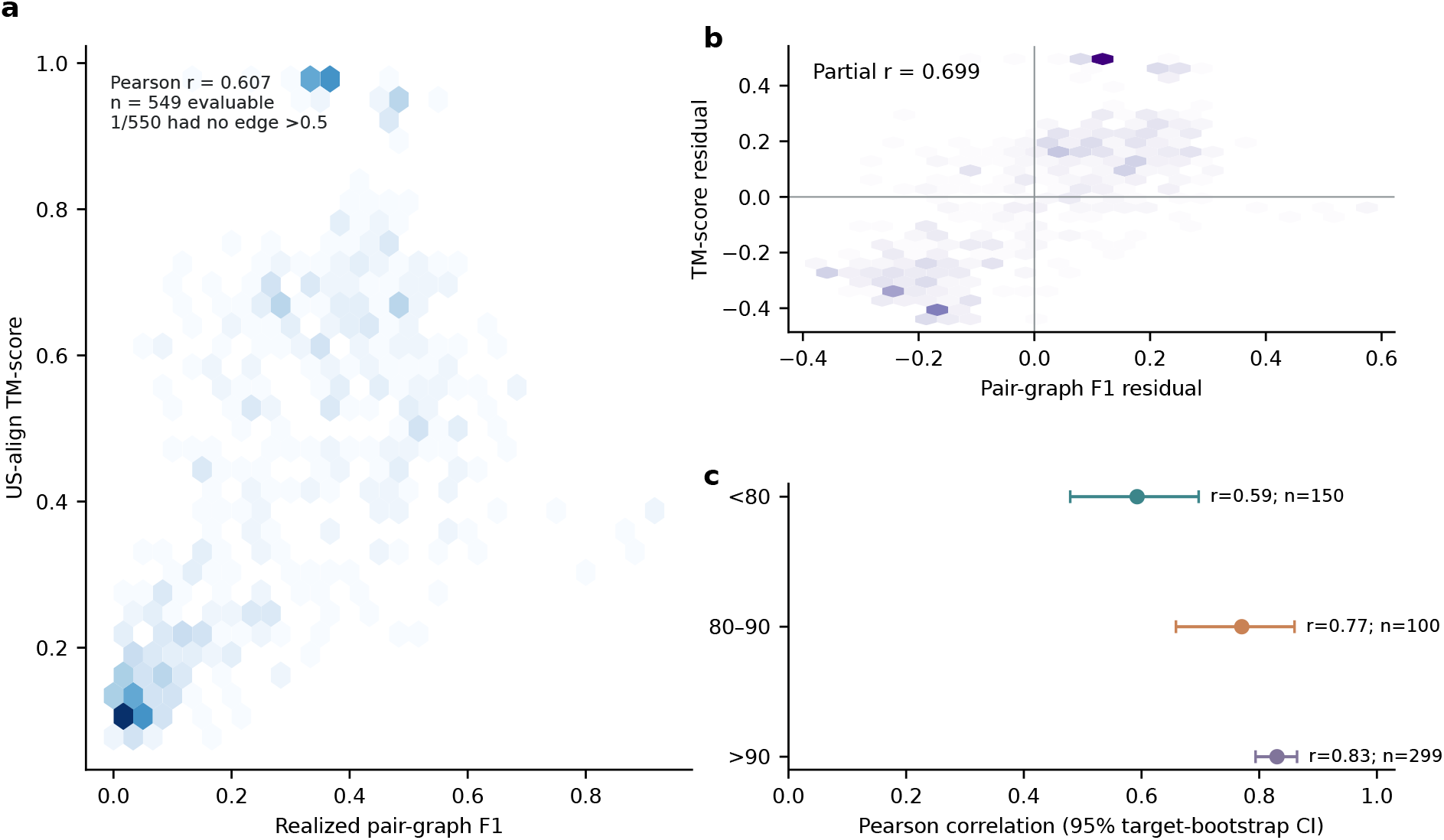
Pair-graph realization is associated with structural compatibility after accounting for RNA length. **a**, Relationship between realized pair-graph F1 and TM-score calculated with US-align across 549 evaluable designs; one additional backbone had no thresholded pair graph because no predicted binary-pair probability *p*_*ij*_ exceeded 0.5. **b**, Residual association after linearly adjusting both variables for chain length. **c**, Pearson correlations within three length strata; intervals are 95% target-bootstrap confidence intervals. The association persists after length adjustment and within strata.

**Extended Data Figure 2.**
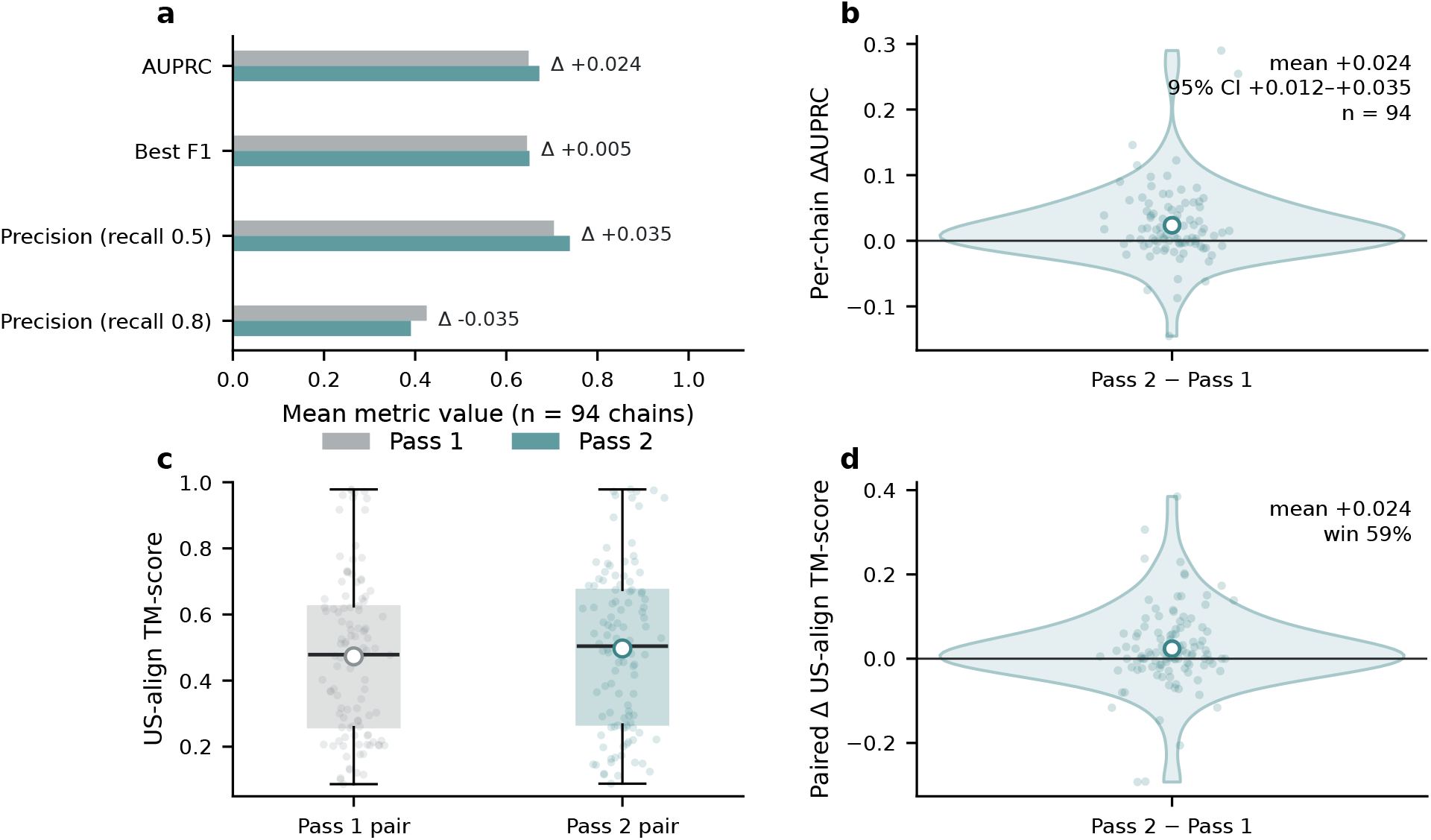
Pair recycling changes pair ranking and downstream sequence decoding. **a**, Mean pair-ranking metrics for Pass 1 and Pass 2 on the 94-chain native benchmark; AUPRC and best F1 increased, while precision at a recall of 0.8 decreased. **b**, Per-chain Pass-2-minus-Pass-1 change in AUPRC, with the mean change and its 95% confidence interval annotated. **c**, Paired TM-scores when the same decoder receives Pass 1 or Pass 2 binary pair-logit inputs while the remaining decoder inputs are fixed; large points denote cohort means. **d**, Distribution of paired TM-score differences.

**Extended Data Figure 3.**
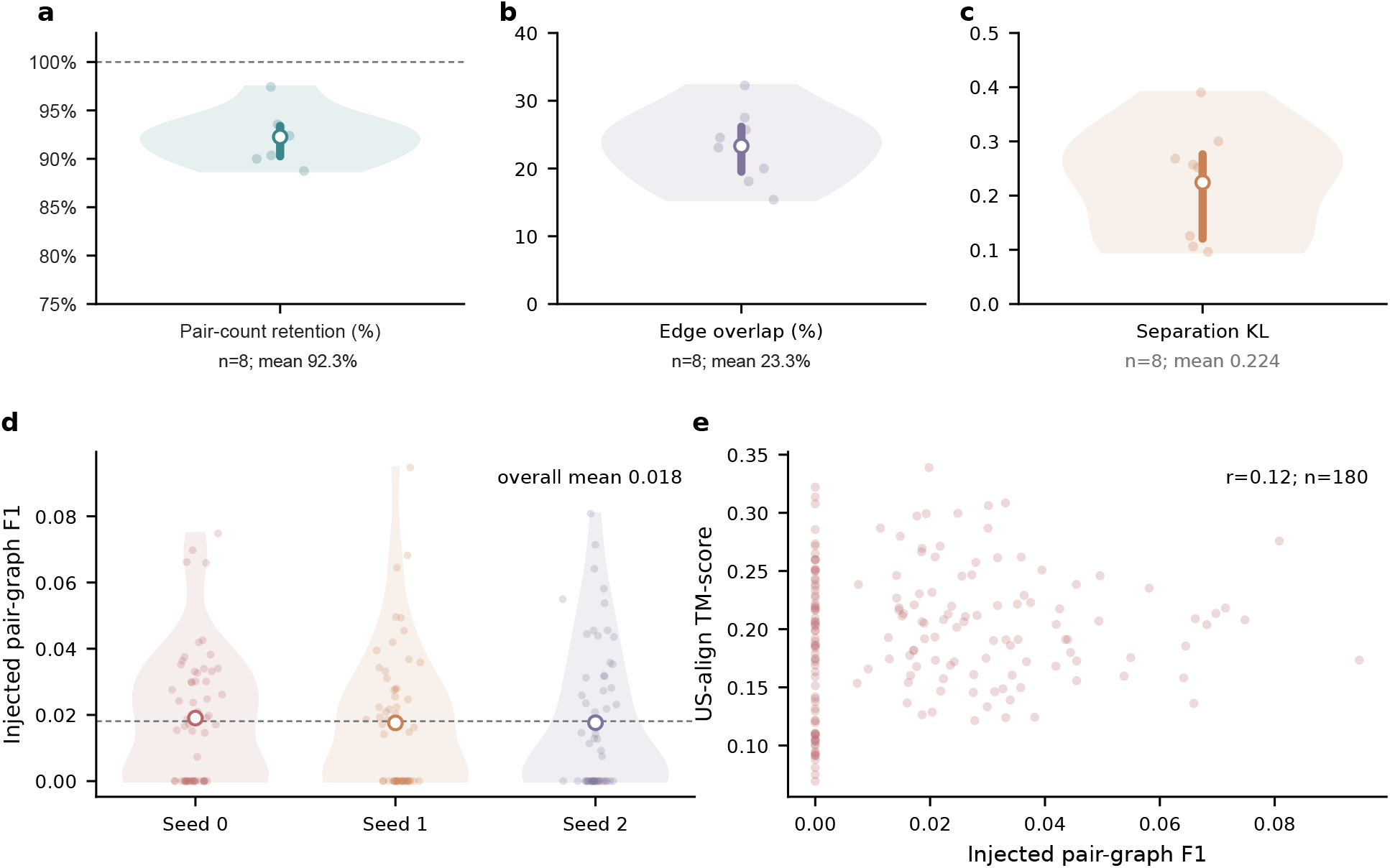
Properties of matched-adversarial rewiring. **a–c**, Construction diagnostics in eight backbones for pair-count retention, residual edge overlap and divergence between original and rewired sequence-separation distributions. Rewired graphs retained, on average, 92.3% of original pair counts while changing residue partners. **d**, F1 between injected adversarial edges and pairs extracted from refolded structures across three seeds. **e**, Relationship between injected-graph F1 and TM-score. Low F1 indicates that injected edges rarely matched refolded pair contacts; the weak association in **e** shows that injected-graph F1 varied little with TM-score. Panels **a–c** and **d**,**e** use separate construction and retention analyses.

**Extended Data Figure 4.**
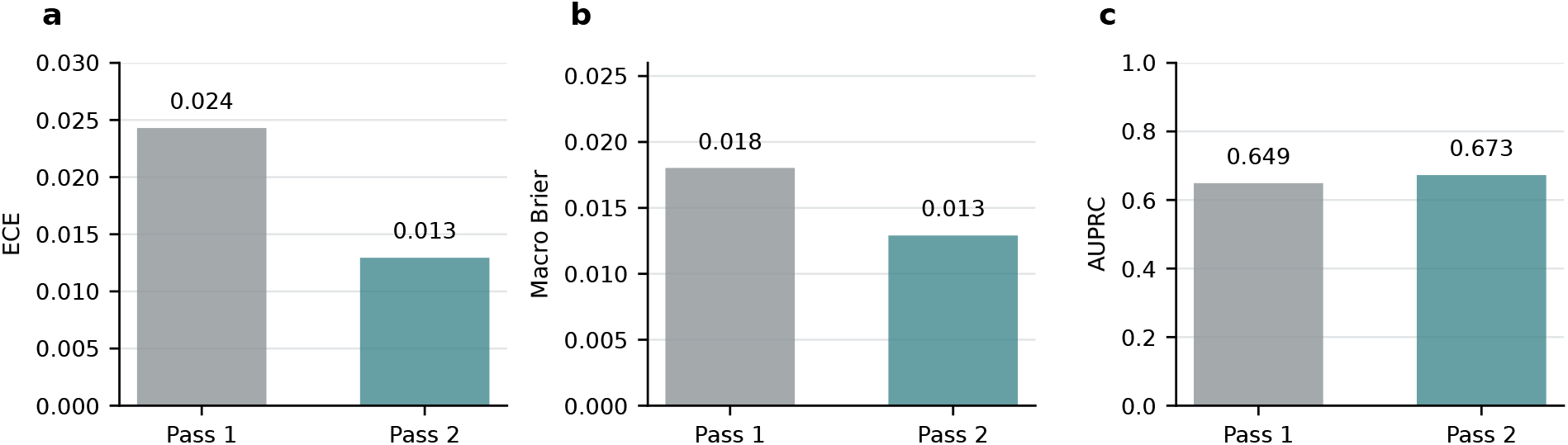
Pair recycling improves probability calibration on the 94-chain native benchmark. First- and second-pass pair probabilities were evaluated against hydrogen-bond-based pair annotations across 377,234 valid undirected residue pairs (distinct from the X3DNA-DSSR-derived canonical reference used for the Fig. 5 sequence-design metrics). **a**, Pooled expected calibration error across all edges. **b**, Target-macro Brier score; the paired Pass-2-minus-Pass-1 comparison and its 95% target-bootstrap confidence interval are reported in Supplementary Table S8b. **c**, Target-macro area under the precision–recall curve. Lower values indicate better calibration for ECE and Brier score, whereas higher AUPRC indicates better pair discrimination. Recycling reduced calibration error and increased AUPRC on this cohort.

**Extended Data Figure 5.**
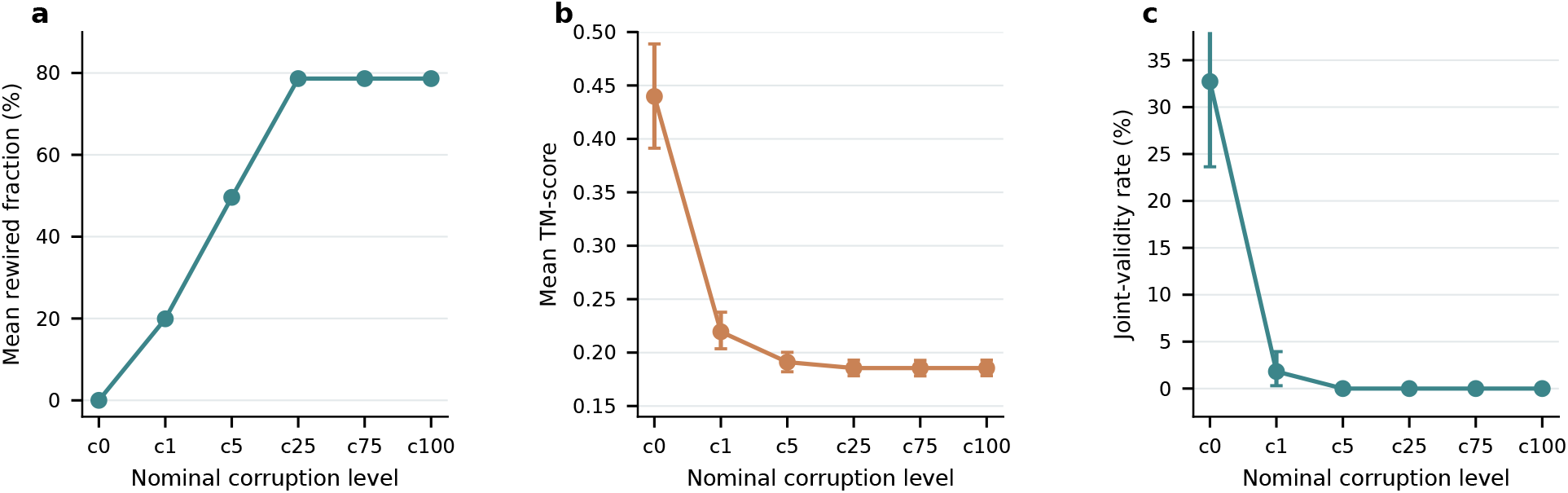
Partial partner rewiring perturbs structural recovery. Results use a separately sampled 110-backbone partner-rewiring corruption-sensitivity cohort and the nominal corruption levels c0, c1, c5, c25, c75 and c100 (cX denotes nominal X% corruption), with three rewiring seeds per level. **a**, Mean achieved rewiring fraction for each nominal level. **b**,**c**, Mean TM-score and joint-validity rate across the nominal corruption levels.

**Extended Data Figure 6.**
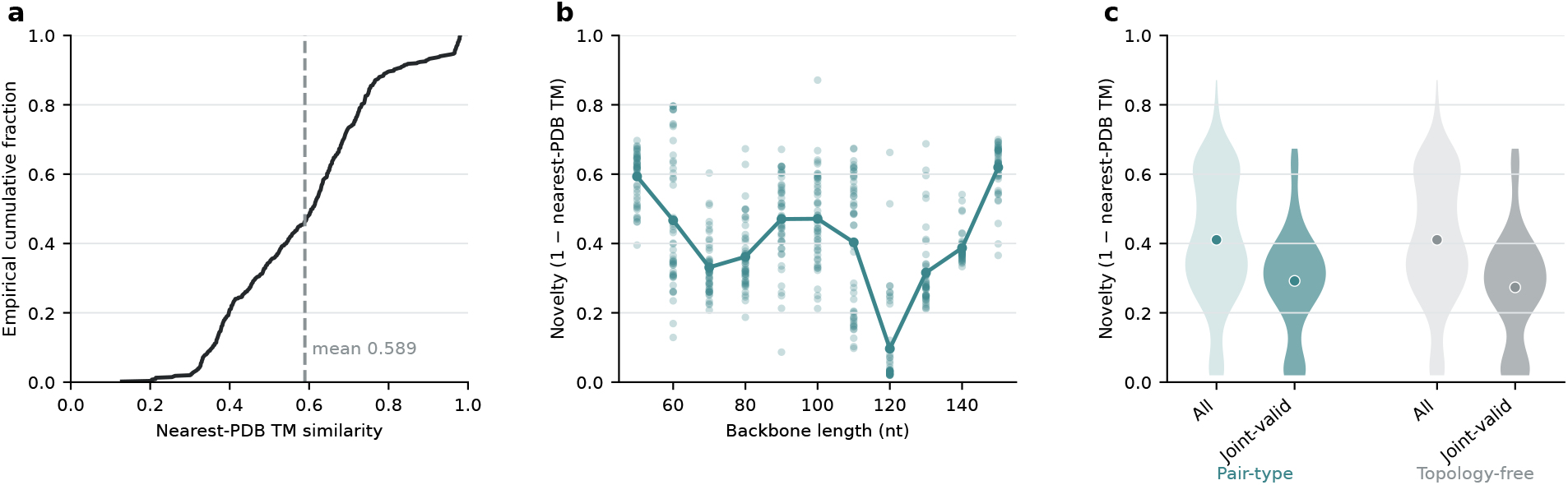
Generated-backbone novelty and validity-associated selection. **a**, Empirical cumulative distribution of nearest-PDB TM similarity for the 550 generated backbones; the dashed line marks the mean of 0.589. **b**, Per-backbone novelty, defined as one minus nearest-PDB TM similarity, across RNA length; the line connects length-wise means. The 120-nt subset showed higher similarity to the reference PDB library than the other length groups. **c**, Novelty distributions before and after joint-validity filtering for pair-type and topology-free decoding. Because these conditions share the same backbone pool, the unfiltered distributions are identical; shifts among joint-valid subsets reflect selection rather than decoder-driven changes in generated geometry.

